# A Long-lived Avatar for Modeling Age-Related Vascular Disease

**DOI:** 10.64898/2026.04.29.721776

**Authors:** Weifeng Qin, Tu N Tran, Yiwei Xiao, Katelyn Ta, Ibrahem Kandel, Ankit Kumar, Rashmi Pandey, Bao Minhngoc A. Dongchau, Vrutant V. Shah, Thi Kim Cuc Nguyen, Kaylee N. Carter, Kristopher W. Brannan, Guangyu Wang, Abhishek Jain, Anahita Mojiri, John P. Cooke

**Affiliations:** Department of Cardiovascular Sciences, Houston Methodist Research Institute, 6670 Bertner Ave, Houston, TX 77030, USA; Department of Biomedical Engineering, Texas A&M University, College Station, Texas 77843, USA

**Keywords:** Vascular avatar, endothelial cells, vascular smooth muscle cells, senescence

## Abstract

**Background:** Current microphysiological models do not support long-term investigations into the chronic effects of vascular risk factors and the development of vascular diseases. Prolonged culture frequently leads to cellular senescence and loss of functional integrity, resulting in variability and inconsistency in modeling chronic vascular responses. Here we aimed to develop and sustain a long-term multicellular human vascular avatar, addressing the critical need for long-term disease modeling and drug testing.

**Methods:** To identify the optimal media for longevity, cell identity and function were assessed by morphology, qPCR, beta-gal staining, ELISA, bulk RNA-seq and single cell RNA-seq analysis. After optimizing the culture media, iPSCs-derived ECs and VSMCs from unaffected and Hutchinson-Gilford Progeria Syndrome (HGPS) donors were grown in Gravitational Lumen Patterning (GLP) Vessel- Chips for 1-6 months to generate a long-lived vascular avatar for the study of vascular aging.

**Results:** Guided by quantitative morphological analyses and bulk RNAseq profiling, we generated a novel optimized culture media VSL (VEGF, SB431542 as a TGF-β inhibitor, low fetal bovine serum) that enhances the long-term health of vascular endothelial cells (ECs). Furthermore, we modified the VSL formulation (mVSL) by modulating 8Br-cAMP, FGF, PDGF, and a cell viability enhancer HMH1015 levels to enhance EC-VSMC (vascular smooth muscle cell) crosstalk and support long-term cellular viability. Subsequently, we maintained and characterized a human vascular avatar with a three-dimensional extracellular matrix environment and 3D vascular architecture for over 180 days. Finally, we demonstrated that this long-lived human vascular avatar enabled modeling vascular aging using iPSC-derived vascular cells from patients with Hutchinson-Gilford Progeria Syndrome (HGPS).

**Conclusions:** We have successfully engineered and maintained a human vascular avatar for over 180 days. The vascular avatar provides a robust platform for modeling disease-associated vascular aging and for evaluating therapeutic strategies targeting chronic vascular disorders.

## Introduction

Vascular dysfunction initiates and contributes to the progression of major cardiovascular diseases^1^. Accordingly, experimental models of vascular dysfunction are useful to understand pathophysiology and to discover therapeutic avenues toward improving cardiovascular health. However, conventional animal models are costly, time-consuming, and often lack translational validity, limiting the relevance of mechanistic studies to human disease. Recent bioengineering advances now enable in vitro reconstruction of microvascular structures that mimic the structure and functions of native vessels^2^. However, less attention has been given to maintaining cellular health and identity during long-term multicellular growth in vessel-on-chip systems.

While animal models can approximate cardiovascular pathophysiology^3,4^, genetic and epigenetic differences from humans limit their translational relevance^5^. Current models often require genetic modification, weeks of specialized diets, extensive animal breeding and handling, and costly monitoring equipment, making them time- and resource-intensive. Moreover, the widespread use of experimental animals poses ethical concerns^6,7^. Due to the limited translational value and ethical concerns associated with animal testing, the FDA has championed in vitro human-based systems, such as organ-on-a-chip technology, to improve the predictive relevance of preclinical drug testing while simultaneously reducing or replacing animal use^8,9^.

Various vessel-on-chip systems have been developed^10–12^, including microfabricated channels, hydrogel-based self-assembled networks, and 3D bioprinted vascular constructs, each aiming to recapitulate physiological flow and vessel architecture^13^. Gravitational Lumen Patterning (GLP) of Vessel-Chips describes the formation of a vascular lumen created by gravity forcing buffer through an extracellular matrix (ECM) that incorporates vascular smooth muscle cells (VSMCs) inside the Vessel-Chips^14,15^. VSMCs growing in the ECM mimic the vascular tunica media, creating a vascular wall, whereas the lumen generated by buffer penetration generates a luminal surface for the culture of an endothelial monolayer. When mature, this platform structurally and physiologically mimics human blood vessels and has potential for vascular biologists and pathologists to study chronic vascular diseases if successfully maintained for extended periods. However, maintaining vascular cell viability over extended periods remains challenging due to cell death, senescence and progressive loss of cellular identity and viability^16,17^.

Many studies have identified individual factors that can enhance endothelial cell (EC) longevity by inhibiting apoptosis through activation of the PI3K/Akt/FoxO signaling pathway (e.g., VEGF) ^18^, as well as by improving mitochondrial activity^19^, regulating metabolism, and reducing inflammation and senescence^18,20^. Additional modulators, such as transforming growth factor-β (TGF-β) inhibitors, have been shown to prevent endothelial-to-mesenchymal transition (EndoMT) induced by oxidative stress and hypoxia^21–23^. Additionally, key regulators such as platelet-derived growth factor (PDGF)^24^, TGF-β^25^, and fibroblast growth factor-2 (FGF2)^26^ promote VSMC proliferation, whereas nitric oxide (NO), cyclic GMP, and AMP-activated protein kinase (AMPK) signaling preserve the contractile, non-synthetic phenotype^27,28^. However, to date, these factors or their combinations have not been successfully employed to maintain a stable vascular avatar suitable for studying chronic diseases.

By leveraging the GLP platform and testing the effects of combinations of longevity factors on cell function and identity, we aimed to develop and sustain a long-term multicellular human vascular avatar, addressing the critical need for long-term disease modeling and drug testing.

## Results

### Identifying Factors that Maintain EC Viability and Identity

To generate a long-term EC culture, we first modified culture media by screening well-known EC viability factors in a 24-well plate. We treated human aortic endothelial cells (HAECs) with nine such factors based on expert opinion of 3 endothelial biologists supplemented by the literature^22,29–37^.These factors included VEGF (10 ng/mL); the TGF-β inhibitor SB 431542(10 µM); low FBS (3%); an H₂S donor, NaSH (1.0 mM); NAD+ precursor (0.3 mM); lower temperature (35 °C); 8-Br-cAMP (10-100 µM); 8-Br-cGMP (10-100 µM); and, the GSK-3 inhibitor CHIR 99021 (10 µM). Based on morphological assessment and quantification of cell area and circularity, we identified three factors that best maintained endothelial viability and morphology: VEGF, SB and low FBS (Figure 1A). Among these factors, VEGF enhanced HAEC survival, but resulted in a densely packed endothelial monolayer with an increase in cell density by comparison to ctrl (Figure 1B). Morphological analysis on day 60 demonstrated that in standard culture media EC area and circularity increased (senescent cells are more circular and have greater area^38,39^), an effect which was attenuated by VEGF. SB also promoted HAEC survival and reduced cell area relative to controls, while generating a less dense monolayer. However, this monolayer assumed a cobblestone morphology that is redolent of the activated endothelium that supports inflammation in vivo at sites of vascular injury^40^ or disturbed flow^1^. Low FBS similarly enhanced HAEC survival, and reduced cell circularity at day 60 (Figure 1C, D). In contrast, cAMP or cGMP produced unexpected negative effects. Neither cAMP nor cGMP supported long-term HAEC survival, and the few cells that persisted exhibited high cell circularity and cell area, typical of senescent ECs (Figure 1A).

**Figure 1.**
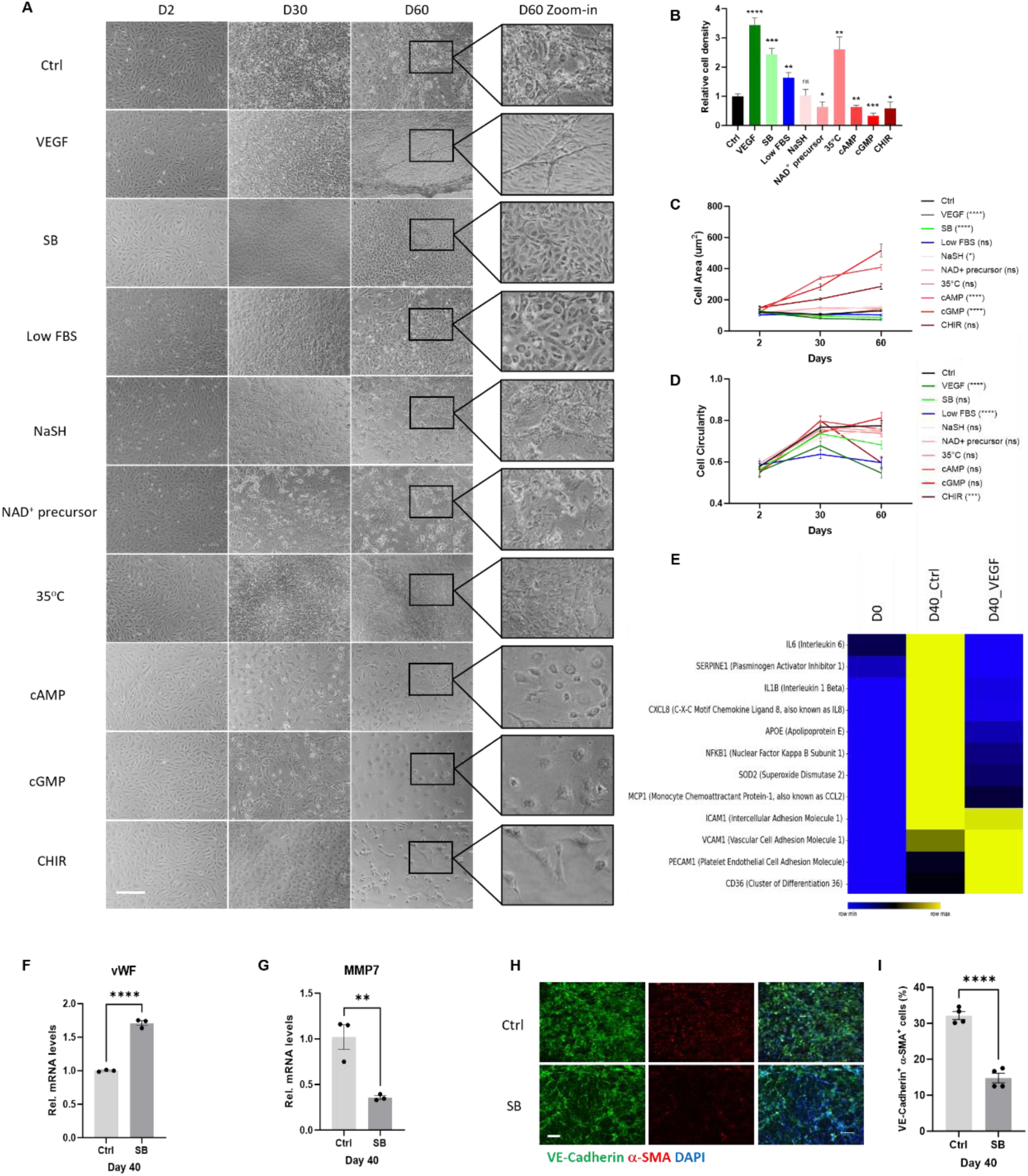
Modifying Culture Conditions to Enhance EC Viability. **A**, Representative images at day 2, 30, and 60 of HAECs treated with different viability factors:Ctrl = standard EC medium (EGM-2MV with 5% FBS); SB = TGF b inhibitor (SB 431542, 10 µM) added to standard EC medium; VEGF= VEGF (10 ng/mL) added to standard EC medium; Low FBS = standard EC medium (3% FBS); NaSH = an H₂S donor, Cat# 161527, 0.3 mM) added to standard EC medium; lower temperature (35 C); cAMP = 100 µM 8-Br-cAMP added to standard EC medium; cGMP = 100 µM 8-Br-cGMP added to standard EC medium; CHIR = GFK-3 inhibitor (CHIR 99021, 10 ng/mL) added to standard EC medium (scale bar: 200 µm). **B**, Quantification of cell density at day 60 from (**A**). Cell density from each group is compared to ctrl at day 60 (n=3). **C,** Quantification of cell area at day 60 from (**A)**. **D**, Quantification of cell circularity at day 60 from (**A).** Cell circularity was assessed as previously described. A cell circularity score of 1 is perfectly circular, whereas a cell circularity score of 0 is perfectly linear. An increase in score indicates greater circularity. Endothelial cells that have higher scores are more circular, and more likely senescent. **E,** analysis of inflammation markers from bulk RNA-seq data in HAECs at day 0 and day 40 treated with or without VEGF reveals that VEGF reduces inflammatory cytokines. **F**, qPCR analysis of EC marker vWF in HAECs treated with or without SB at day 40 reveals that SB preserves EC identity (n=3). Each dot represents one technical repeat. **G**, qPCR analysis of fibroblast marker MMP7 in HAECs treated with or without SB at day 40 reveals that SB reduces expression of a fibroblast marker (n=3). Each dot represents one technical repeat. **H**, Representative images of immunofluorescence staining of EC marker VE-Cadherin and fibroblast marker a-SMA in HAECs treated with or without SB at day 40 are shown (scale bar: 200 µm, n=3). **I**, Quantification of the percentage of VE-Cadherin and a-SMA double positive cells from (**H**) is consistent with an effect of SB to prevent EndoMT. Each dot represents one single field from at least three technical repeats. Data between two groups were analyzed by Student’s t-test. Data between multiple groups are analyzed by one-way ANOVA. Results are considered statistically significant with P<0.05(*), P<0.01(**), P<0.001(***), and P<0.0001(****).

To confirm the morphological observations, heatmap analysis of inflammatory markers further revealed that VEGF partially attenuated inflammatory gene expression (Figure 1E). In contrast, the cAMP- and cGMP-treated groups clustered distantly from all other conditions, with transcriptional signatures that were consistent with the morphological evidence of senescence (Supplementary Figure 1A).Since SB is a well-known inhibitor of the TGF-*β* signaling pathway and EndoMT^41^, we examined the extent to which SB maintained HAEC identity. Consistent with our morphometric observations and our hypothesis, qPCR analysis revealed that by comparison to control SB up-regulated EC marker von Willebrand factor (vWF) and down-regulated fibroblast marker matrix metalloproteinase-7 (MMP7) (Figure 1F, G), suggesting that SB prevents HAECs from differentiating into fibroblast-like cells. Supporting this finding, immunofluorescence staining showed that SB reduced the percentage of VE-Cadherin and α-SMA double positive cells (Figure 1H, I), directly demonstrating that TGF-*β* inhibition preserves HAEC identity by suppressing EndoMT.

Both VEGF and TGF-*β* inhibition supported aspects of HAEC health; however, each treatment also revealed some deficiencies. We performed bulk RNA-sequencing data to assess the transcriptional profiles of cells in the various treatment groups. Principal Component Analysis (PCA) of HAECs treated with VEGF demonstrated that the transcriptional profile at a later time point (day 40) was distant from that at day 0, indicating VEGF was not able to preserve the baseline cell transcriptional profile (Figure S1A). Furthermore, VEGF promoted robust early HAEC maintenance but eventually activated neuronal-associated pathways by day 40, suggesting dynamic transcriptional shifts during extended exposure to VEGF (Figure S1B). TGF-*β* inhibition preserved HAEC identity yet was associated with modest increases in inflammatory signals over time, consistent with the morphological alterations (Figure S1C, D).

### Identifying Combinations That Enhance EC Longevity

Based on these findings, we hypothesized that a combinatorial approach integrating all three longevity factors into the culture medium would balance the deficiencies of the factors when given in isolation. To test this, we supplemented complete EC medium with VEGF (10 ng/mL), SB (10 uM) and low FBS (3%) (hereafter referred to as VSL). Because our long-term goal is to apply this strategy to patient-derived samples, we evaluated VSL in ECs differentiated from induced pluripotent stem cells (iPSC). The iPSC-derived ECs provide a relevant human model for personalized vascular studies^42,43^. The iPSC-ECs were cultured under different conditions and monitored for 40 days. Regular morphometric analyses of cell area and circularity were performed to quantify the effects of VSL during the time. The results showed that VSL improved iPSC-EC morphology, reducing cell circularity more effectively than either VEGF or SB alone over a 25-day culture period (Figure 2A-C). To further validate these observations, we performed transcriptional profiling with UMAP analysis on iPSC-ECs collected at day 40 following treatment with VEGF alone, VSL, or VSL supplemented with cAMP and/or cGMP. In some cases, we also added a prophylactic antibiotic (Lonza Walkersville MycoZap, 0.2%) against mycoplasma for these long-term cultures. The antibiotic (AB) did not substantially alter the UMAP analysis, and was therefore used in all subsequent long-term studies to avoid mycoplasma contamination. The unbiased UMAP analysis also revealed that VSL, regardless of additional supplementation, successfully restored the transcriptional profile of iPSC-ECs at day 40 to closely resemble that of day 0 (Figure 2D). Thus, the transcriptional studies supported the use of VSL to maintain long-term iPSC-EC viability and longevity.

**Figure 2.**
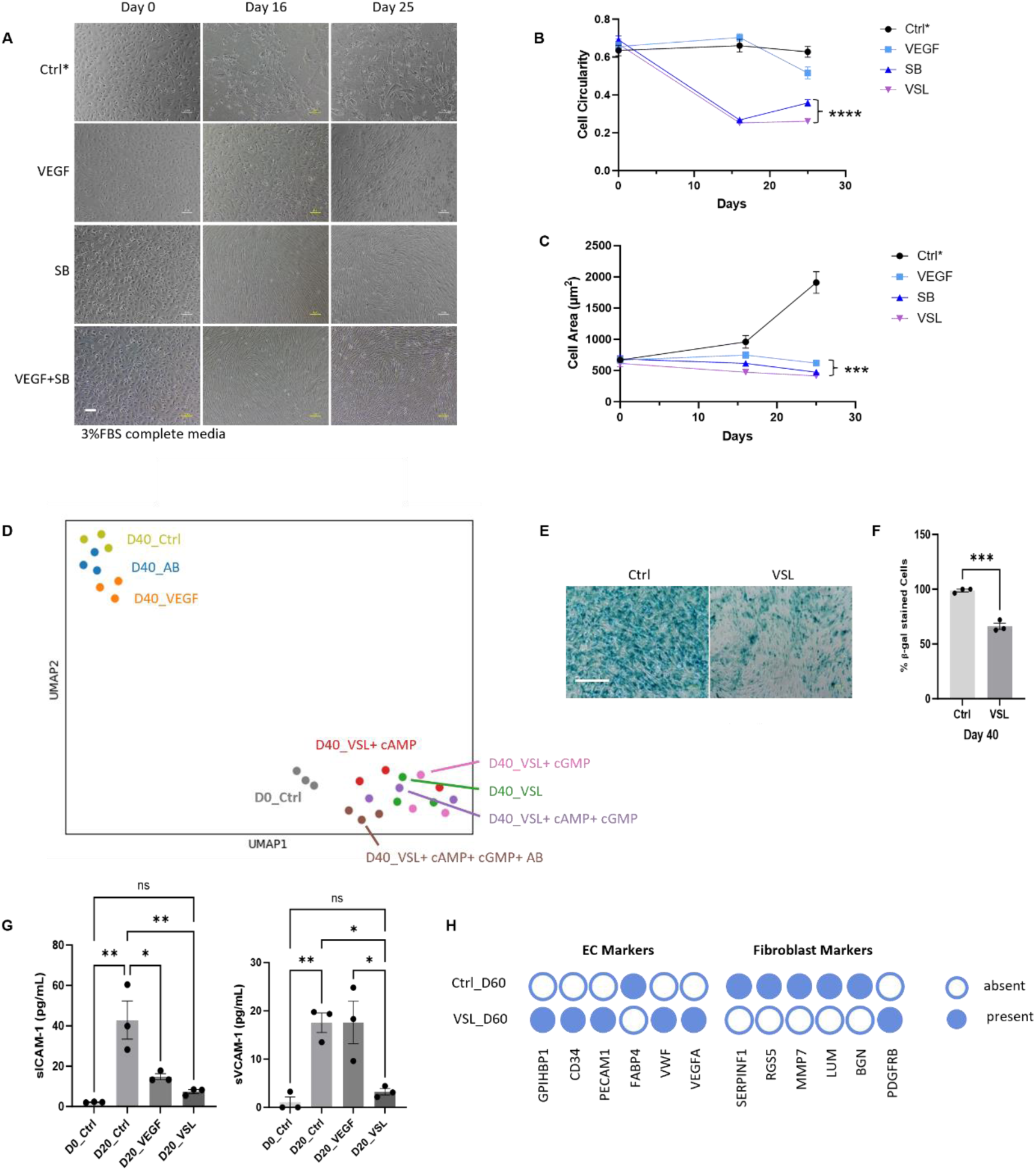
Further enhancement of EC viability by combinatorial treatment. **A**, Representative images of iPSC-ECs in low serum (3%) and treated with VEGF, SB, or VEGF+SB (VSL) at the indicated time points reveals that VSL further improves EC morphology. The above treatments were performed using EC medium (Lonza, EGM-2MV) with 3% FBS. Ctrl* = EC medium with 3% FBS; VEGF = VEGF (10 ng/mL) added to EC medium with 3% FBS; SB = SB (SB 431542, 10 µM) added to EC medium with 3% FBS; VEGF + SB = VEGF (10 ng/mL) and SB (SB 431542, 10 µM) added to EC medium with 3% FBS, referred to as VSL. (scale bar: 100 µm) **B**, Quantification of cell circularity from (**A**) reveals that VSL reduces cell circularity. **C**, Quantification of cell area from (**A**) reveals that VSL reduces cell area. **D**, UMAP analysis of iPSC-ECs treated with different viability factors or combinatorial treatments of viability factors at day 40 shows that VSL restores the transcriptional profile of HAECs at day 40 to that at day 0 (n=3). The prophylactic antibiotic (Lonza Walkersville MycoZap, 0.2%; AB) was used to prevent mycoplasma contamination. D0_Ctrl = iPSC-ECs at day 0; D40_Ctrl = iPSC-ECs treated with 3%FBS EC medium (Lonza, EGM-2MV) for 40 days; D40_VEGF = iPSC-ECs treated with the 3%FBS EC medium with the addition of VEGF (10 ng/mL) for 40 days; D40_AB = iPSC-ECs treated with 3%FBS EC medium + prophylactic antibiotic (Lonza Walkersville MycoZap, 0.2%; AB) for 40 days; D40_VSL = iPSC-ECs treated with VSL for 40 days; D40_VSL+cAMP = iPSC-ECs treated with VSL and cAMP (10 µM) for 40 days; D40_VSL+cGMP = iPSC-ECs treated with VSL and cGMP (10 µM) for 40 days; D40_VSL+Camp + cGMP = iPSC-ECs treated with VSL, cAMP (10 µM), and cGMP (10 µM) for 40 days; D40_VSL + cAMP + cGMP + AB = iPSC-ECs treated with VSL, cAMP (10 µM), cGMP (10 µM) and AB (Lonza Walkersville MycoZap, 0.2%;) for 40 days. **E**, Representative images of β-gal staining for iPSC-ECs treated with or without VSL at day 40 are shown (scale bar: 200 µm). **F**, Quantification of **(E)** shows that VSL reduces the percentage of β-gal positive cells (n-=3). Each dot represents one single field. **G**, The expression of sICAM-1 and sVCAM-1 in culture media from iPSC-ECs treated with VEGF alone or VSL at day 0 or day 20 detected by ELISA assay shows that VSL effectively suppresses inflammatory cytokines (n=3). D0_Ctrl = iPSC-ECs at day 0; D20_ Ctrl = iPSC-ECs treated with standard EC medium (Lonza, EGM-2MV) for 20 days; D20_VEGF = iPSC-ECs treated with standard EC medium and addition of VEGF (10 ng/mL) for 20 days; D20_VSL = iPSC-ECs treated with VSL for 20 days. Each dot represents one technical repeat. **H**, Heatmap analysis of EC marker genes and fibroblast marker genes expressed in iPSC-ECs treated with or without VSL at day 60 reveals that VSL preserves EC identity during long-term culture. Each group includes triplicate in this analysis. Ctrl_D60 = iPSC-ECs treated with 3%FBS EC medium (Lonza, EGM-2MV) for 60 days. VSL_D60 = iPSC-ECs treated with VSL. Data between two groups were analyzed by Student’s t-test. Data between multiple groups are analyzed by one-way ANOVA. Results are considered statistically significant with P<0.05(*), P<0.01(**), P<0.001(***), and P<0.0001(****).

To confirm the effect of VSL to maintain viability and identity of long-term iPSC-EC culture, we performed a heatmap analysis of the transcriptional data with a focus on genes regulating the established hallmarks of aging^44^. The analysis revealed that VSL inhibited the expression of genes associated with cellular senescence and inflammation; maintained normal epigenetic modifications, suppressed macro-autophagy, supported normal intercellular communication; and sustained nutrient sensing, proteostasis, as well as mitochondrial and telomere function (Figure S2A-I). Consistent with these findings, by comparison to the group, VSL reduced the percentage of β-galactosidase (β-gal) positive iPSC-ECs day 40 (Figure 2E, F). We observed increased release of sICAM-1 and sVCAM-1 into the standard EC media, as measured by ELISA at day 20 of iPSC-EC culture. By contrast, VSL effectively reduced the levels of these soluble adhesion molecules (Figure 2G). Notably, this suppression of sVCAM and sICAM by VSL medium was more pronounced than that achieved by VEGF alone (Figure 2G). Furthermore, heatmap analysis of EC identity genes in iPSC-ECs- treated with or without VSL showed that VSL maintained key endothelial markers (GPIHBP1, CD34, CD31, vWF and VEGFA) while inhibiting the expression of fibroblast-associated genes (SERPINF1, RGS5, MMP7, LUM, and BGN). (Figure 2H). Our findings suggest that VSL supports iPSC-EC longevity, by suppressing senescence and EndoMT.

### Establishing a Long-Term 3D Endothelial Monolayer within a Vessel-Chip

Subsequently, we seeded either HAECs or iPSC-ECs into a 3D microfluidic vessel-chip and cultured them in VSL or standard EC medium. Monthly histological analyses showed that HAECs in standard medium remained viable for only 3 months, whereas HAECs in VSL remained viable for over 6 months (Figure 3A), far exceeding the duration of EC viability reported for conventional 3D models (Fig. 5f). We do not know what is the upper bound for HAECs under these conditions as we terminated the experiment at 210 days. Quantitative morphometric analysis further confirmed that VSL maintained normal cell circularity and area (Figure 3B, C).

**Figure 3.**
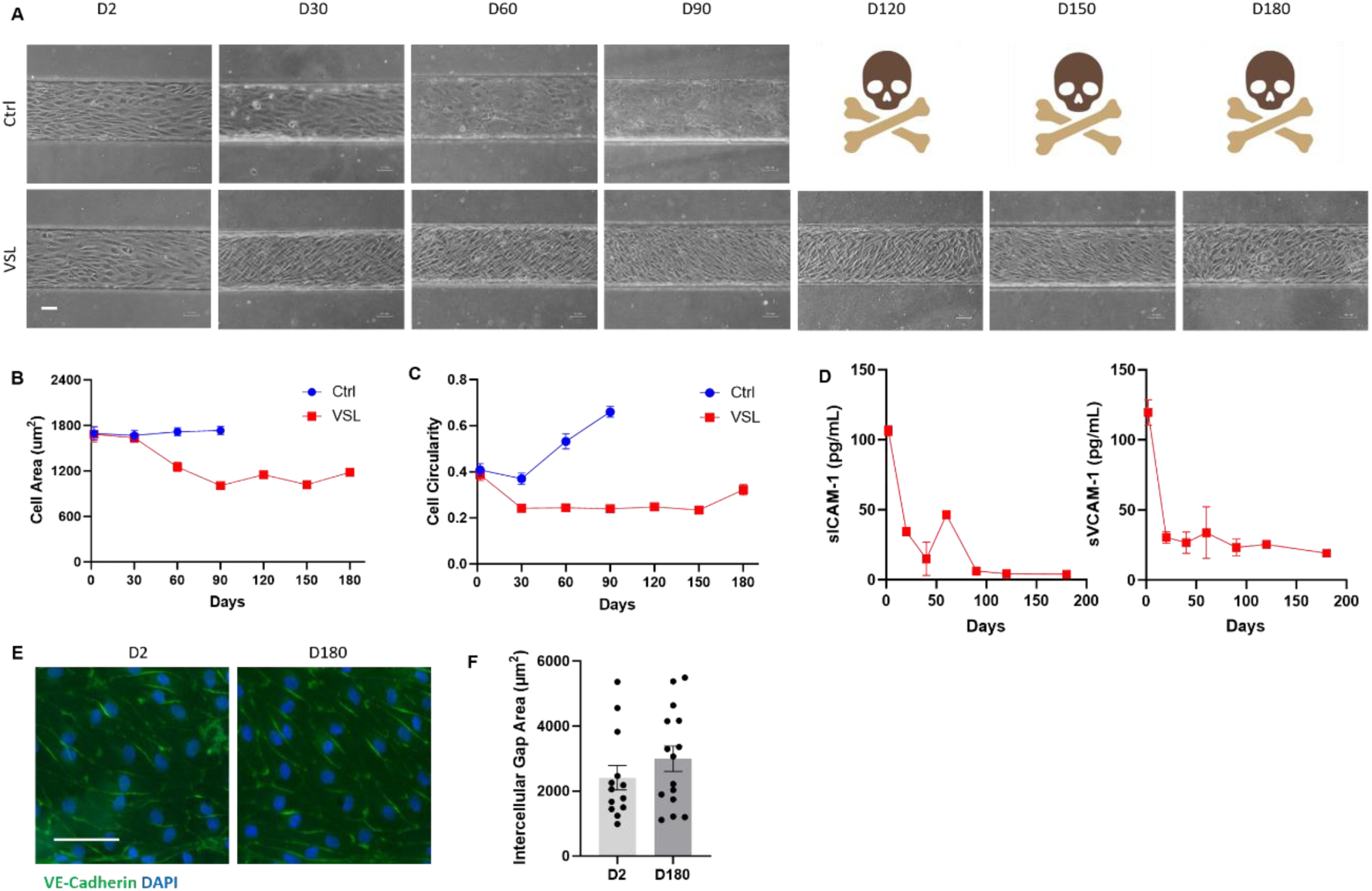
Maintaining a human EC monolayer in vessel-chip for over 180 days by VSL. **A**, Representative images of HAECs show that VSL successfully maintains an intact EC monolayer and cellular morphology for over 180 days, whereas cells cultured in control medium (EGM-2MV with 5%) lose viability and fail to survive beyond 90 days (scale bar: 100 µm). **B**, Quantification by cell area in panel (**A)** shows that throughout the 180-day culture, VSL keeps the cell area lower than that of cells exposed to standard media (Ctrl). Over 30 cells were counted in each time point from each group. **C**, Quantification by cell area in panel (**A**) shows that throughout the 180-day culture, VSL maintains cell circularity at a normal level compared to that of cells exposed to standard media (Ctrl). Over 30 cells were counted in each time point from each group. **D**, VSL reduces the expression of sICAM-1 and sVCAM-1 in the EC monolayer compared to day 0 throughout the 180-day culture. (n=3). **E**, Immunofluorescence staining for the EC monolayer shows that EC marker VE-Cadherin is preserved by VSL at day 180 days. (scale bar: 125 µm). **F**, Quantification of (**E**) using intercellular gap area indicates that cell permeability does not increase at day 180 compared to that at day 2.

**Figure 4.**
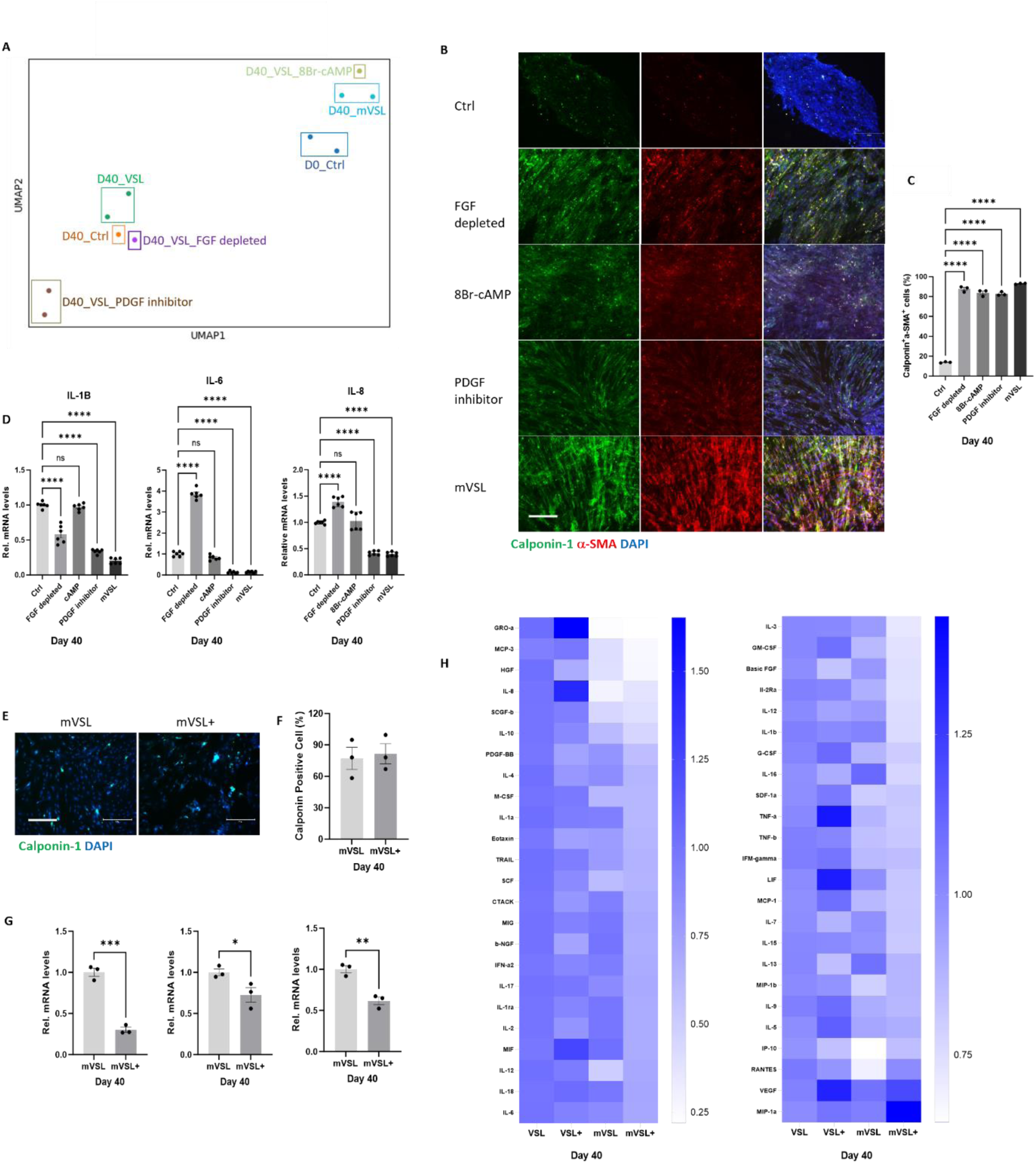
Optimizing Culture Conditions to Enhance VSMC Identityand Quiescence. **A**, Unbiased UMAP analysis shows that transcriptional profile of VSMCs treated with mVSL is the closest to that of cells at day 0. **B**, Immunofluorescence staining reveals that mVSL is effective in preserving VSMC contractile markers Calponin-1 and α-SMA at day 40. Ctrl = VSL; FGF depleted= VSL with FGF depletion; 8Br-cAMP = 2 µM 8Br-cAMP added to VSL; PDGF inhibitor = PDGF inhibitor (AG 1295, 10µM) added to VSL; mVSL = VSL with PDGF inhibitor (AG1295 10 μM), 8Br-cAMP (2 μM), low serum (1% FBS), and depletion of FGF. (scale bar: 300 µm). **C**, Quantification of (**B**) confirms that mVSL is effective in preserving VSMC contractile markers Calponin-1 and α-SMA at day 40. **D**, qPCR analysis reveals that mVSL effectively reduces inflammatory markers IL-1b, IL-6, and IL-8. at day 40. Ctrl = VSL, FGF depleted= VSL with FGF depletion, 8Br-cAMP = 2 µM 8Br-cAMP added to VSL, PDGF inhibitor = PDGF inhibitor (AG 1295, 10µM) added to VSL, mVSL (VSL with PDGF inhibitor (AG1295 10 μM), 8Br-cAMP (2 μM), low serum (1% FBS) with depletion of FGF). **E**, Immunofluorescence staining shows cells that are Calponin-1 positive at day 40 in VSMCs treated with mVSL or mVSL+ (mVSL+ is mVSL medium to which HMH1015 (v/v 10%; a cell viability factor) is added (scale bar: 300 µm). **F**, Quantification of (**E**) shows that mVSL+ does not reduce Calponin-1 positive cells. **G**, qPCR analysis shows that mVSL+ further reduces inflammation markers IL-1b, IL-6, and IL-8 at day 40. **H**, Bio-plex analysis shows that mVSL+ reduces most of the inflammatory markers detected in the culture media at day 40. VSL+ = HMH1015 (v/v 10%) added to VSL; mVSL+= HMH1015 (v/v 10%) added to mVSL.

**Figure 5.**
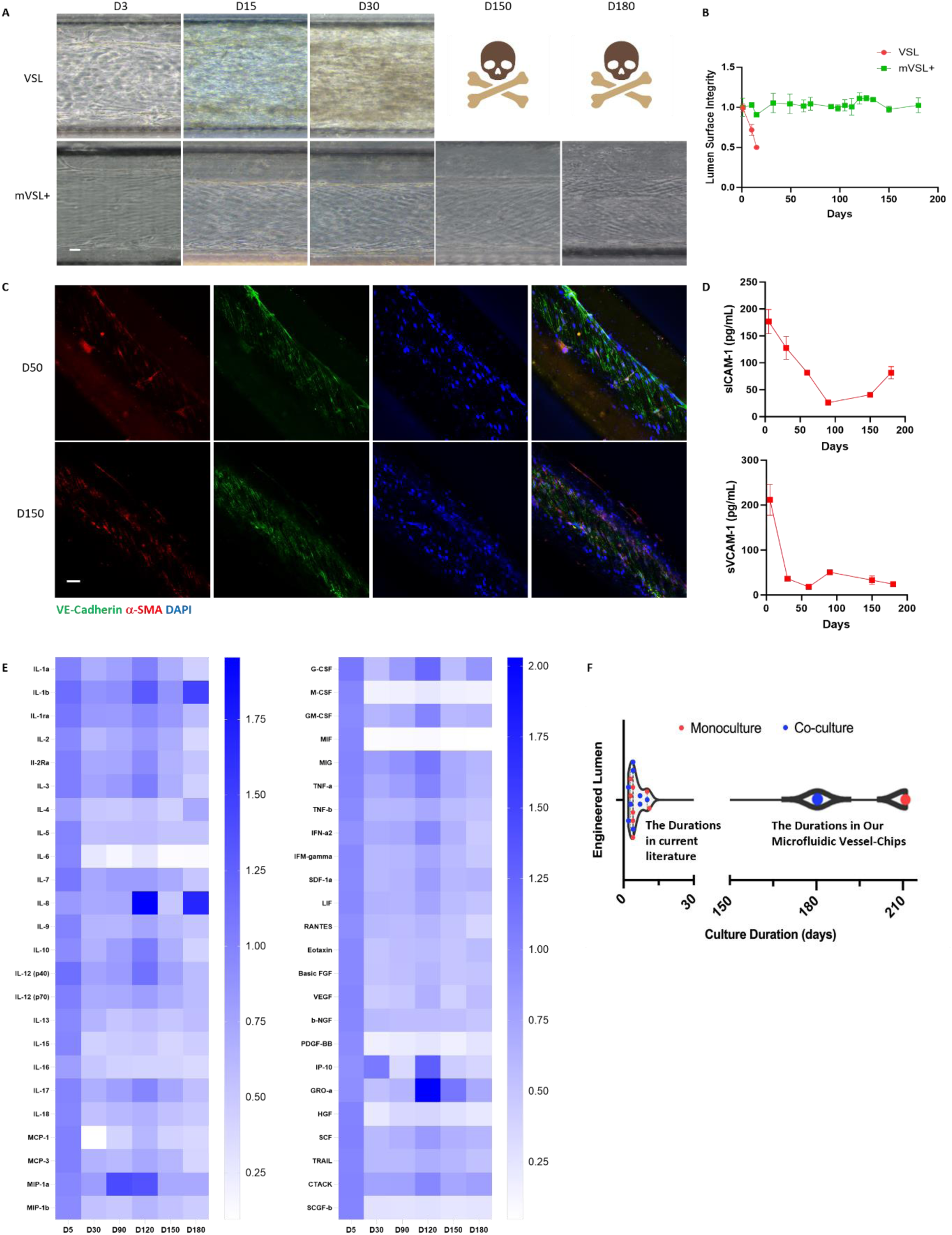
Maintaining a human vascular avatar for over 180 days by mVSL+. **A**, mVSL+ maintains co-culture of human iPSC-ECs and iPSC-VSMCs for over 180 days, with endothelial cells aligning with the lumen direction throughout the 180-day culture (scale bar: 40 µm). **B**, Quantification of (**A**) shows that mVSL+ preserves lumen surface integrity throughout 180-day culture. **C,** Immunofluorescence staining of VE-Cadherin and α-SMA in the avatars at the indicated time points (scale bar: 40 µm). **D**, mVSL+ maintains the expression of sICAM-1 and sVCAM-1 in the vascular avatars at a relatively low level compared to day 5 throughout the 180-day culture. (n=3). **E**, Bio-plex analysis shows that mVSL+ maintains the vascular avatars at a low inflammatory state compared to day 5 throughout the 180-day culture**. F**, The duration of prior 3D models of vascular lumens using monoculture (in red) or co-culture (in blue). Each dot represents the duration noted in one paper. 17 papers are cited in total. The large red dot indicates our monoculture duration, and the large blue dot indicates our co-culture duration.

Furthermore, sVCAM-1 and sICAM-1 levels both decreased over the 180-day culture period in the vessel-chip compared to day 2 (Figure 3D). Immunofluorescence staining of VE-Cadherin revealed that structural junction integrity, quantified by intercellular gap area as previously described^11,45^, was comparable at day 180 to that at day 2 (Figure 3E, F), indicating sustained endothelial barrier integrity by VSL. Consistent with our studies using HAECs, we also maintained a vessel-chip lined with iPSC-ECs for over 180 days. The iPSC-ECs exhibited a normal EC morphology at day 80 (Figure S3A, B), although cell density gradually declined by days 130 and 180 (Figure S3C).

### Identifying Culture Conditions for Long-Term VSMC Viability

Having developed conditions that promote long-term viability of the EC monolayer, we now intended to determine if such conditions would also provide for long-term maintenance of an EC-VSMC co-culture. Unexpectedly, we found that the VSL media was not optimal for EC-VSMC co-culture within the GLP Vessel-Chip primarily due to VSMC over-proliferation that disrupted lumen surface integrity (Figure S4). To enable long-term co-culture, we next sought to further modify the culture conditions to maintain VSMC quiescence.

Major proliferative factors for VSMCs are PDGF and FGF^46,47^ ^24^ ^26^; whereas cAMP is an inhibitor of VSMC proliferation^48^ and migration^49^. In 2D culture, we tested several modified VSL (mVSL) formulations incorporating an inhibitor of PDGF, or depletion of FGF, or depletion of serum, each of which partially suppressed VSMC proliferation over 40 days, as quantified by DAPI-positive cell counts (Figure S5). Ultimately, the combinatorial treatment containing a PDGF inhibitor (AG1295 10 μM), 8Br-cAMP (2 μM), low serum without addition of FGF (termed mVSL) was the most effective. Morphometric analysis showed that VSMCs under mVSL conditions adopted an elongated morphology characteristic of quiescent cells^31^.

Bulk RNA-seq analysis of samples collected at day 40 revealed that the transcriptional profiles of VSMCs treated with mVSL or VSL+8Br-cAMP closely resembled those at day 0, whereas other treatment groups diverged substantially. This finding indicates that mVSL or VSL+8Br-cAMP are the most effective conditions for preserving the VSMC transcriptional program during long-term culture (Figure 4A). Furthermore, immunofluorescence staining revealed that the percentage of VSMC contractile markers Calponin-1 and α-SMA double positive cells increased in each of the modifications of VSL compared to control group (Figure 4B, C). Notably, VSL+PDGF inhibitor or mVSL stood as the most effective treatment in decreasing inflammatory markers, IL-1b, IL-6, and IL-8 revealed by qPCR analysis (Figure 4D). These data collectively support that mVSL is the most effective in sustaining VSMC viability during long-term culture.

While others have suggested 10% to 20% of FBS for VSMC culture^31^, our morphological observations showed that complete serum withdrawal still supported VSMC viability for over 100 days (Figure S6E, F). In contrast ECs failed to survive long-term without FBS supplementation (Figure S6A-D). Our results indicate that 1% or 3% FBS, in the mVSL formulation provides optimal conditions for maintaining both EC and VSMC viability and quiescence. This low-serum condition was more effective in preserving the VSMC contractile phenotype and reducing inflammation, as evidenced by an increased proportion of Calponin-1 and α-SMA double positive cells (Figure S6G, H). Additionally, IL-1b expression was lower under 1% FBS compared to 3% FBS (Figure S6I).

We further optimized our culture conditions by supplementing mVSL with the small molecule HMH1015 (hereafter mVSL+) for long-term VSMC maintenance. Using mVSL+ we found that the percentage of Calponin-1 positive cells remained stable (Figure 4E, F), while the inflammatory cytokine markers, IL-1b, IL-6, and IL-8, were reduced by comparison to mVSL (Figure 4G). Notably, a comprehensive bio-plex analysis showed that mVSL+ reduced most inflammatory markers detected in the culture medium compared to VSL, VSL+, or mVSL (Figure 4H).

In conclusion, through systematic optimization of culture conditions, we established mVSL+ as an effective medium that supports long-term EC-VSMC co-culture while minimizing inflammation and preserving VSMC contractile phenotype.

### The Novel mVSL+ Media Maintains a Long-lived Human Vascular Avatar

Having optimized the culture medium for both ECs and VSMCs, we next established a co-culture of iPSC-derived ECs and VSMCs within the GLP Vessel-Chips. The optimized mVSL+ medium facilitated the persistence of a human vascular avatar for over 180 days (Figure 5A), exceeding existing in vitro co-culture models, which last up to 10 days (Figure 5F). Remarkably, lumen surface integrity was preserved throughout the 180-day culture, whereas cultures maintained in VSL alone showed luminal surface disruption by day 30 (Figure 5B). Immunofluorescence staining confirmed the presence of VE-Cadherin positive ECs and α-SMA positive VSMCs within the vascular avatar at both day 50 and day 150 (Figure 5C). Furthermore, ELISA analysis showed that sVCAM-1 and sICAM-1 levels fell during the 180-day period compared to day 2 (Figure 5D),which could be due to maturation of the ECs at the later time point^50^. Consistently, a comprehensive bio-plex cytokine analysis showed that most inflammatory cytokines remained at relatively low levels throughout the culture period (Figure 5E).

### Single Cell Transcriptional Signatures Validate the Long-Term Vascular Avatar

To comprehensively characterize the cellular composition of our long-term vascular avatar, we collected cells from the devices at day 0 and day 60 for scRNAseq. The scRNAseq data at day 60 revealed that the co-culture had maintained cell type specificity, sustaining two distinct populations (EC and VSMC populations; Figure 6A, C). Comparative trajectory analysis of gene expression profiles revealed that by day 60 1) the proliferating VSMCs were no longer present, suggesting maturation of the co-culture with quiescent VSMCs and 2) arterial like ECs had transitioned toward a capillary/lymphatic-like phenotype (Figure 6A, B, C). These phenotypic shifts likely reflect adaptive transcriptional reprogramming driven by EC-VSMC crosstalk in low shear stress conditions, mimicking a microvascular microenvironment^51–53^. Additionally, cell cycle regulators CDKN1A, CDKN2A, TP53, and RB1 did not change from day 60 to day 0 (Figure 6D), suggesting that the vascular avatars did not become senescent during long-term culture.

**Figure 6.**
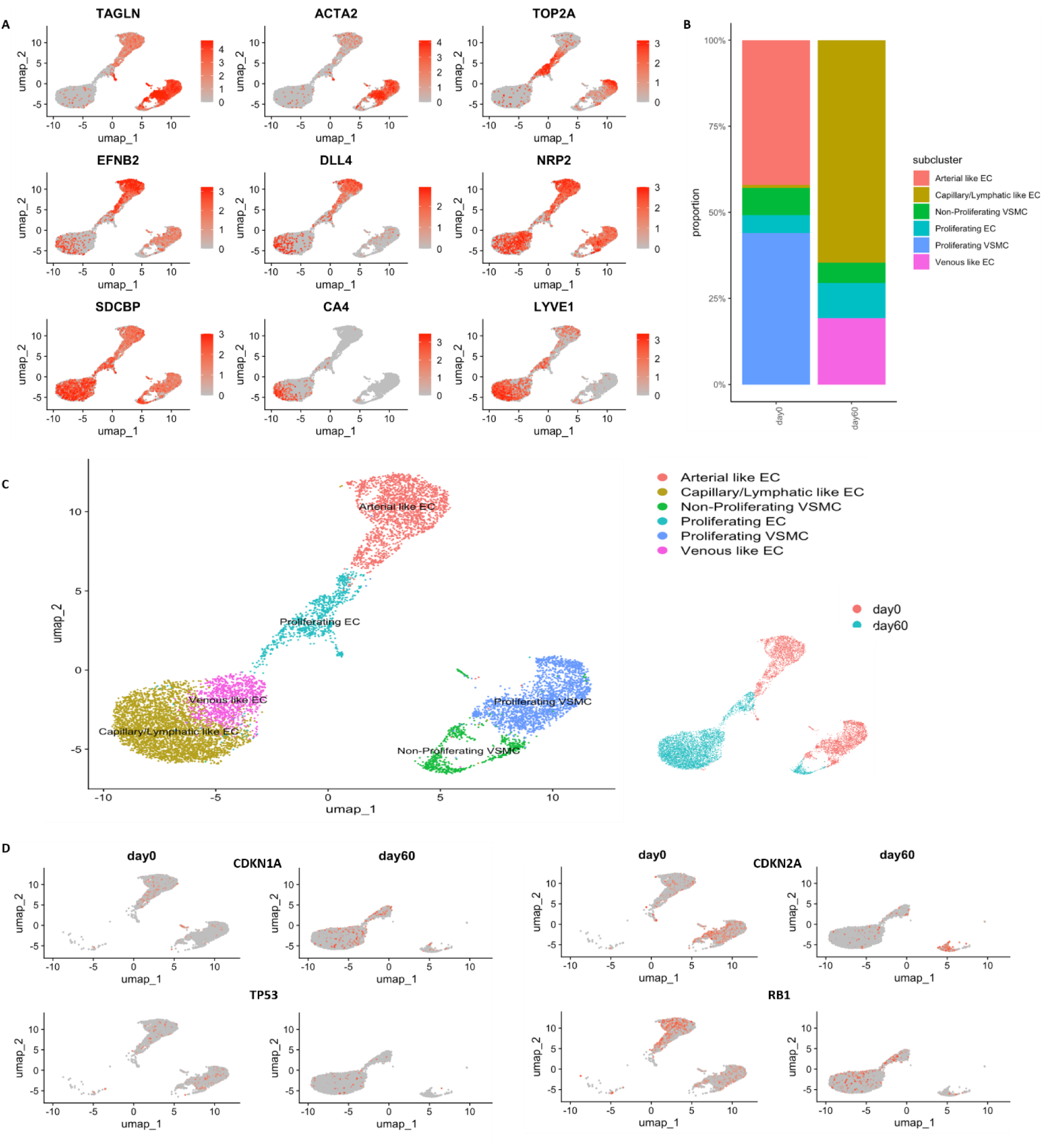
Single cell analysis of the long-term human vascular avatar. **A**, Representative marker genes of different vascular lineage cell types are shown. **B**, The percentages of different vascular lineage cell types from the vascular avatars at day 0 and day 60 are shown. **C**, UMAP plots of all vascular cell populations detected at the vascular avatars at day 0 and 60. **D**, Cell cycle genes CDKN1A, CDKN2A, TP53, and RB1 are shown in the vascular avatars at day 0 and day 60.

### Modelling Vascular Aging in the Avatar

To evaluate the utility of the avatar for modeling vascular aging, we introduced ECs and VSMCs derived from Hutchinson-Gilford Progeria Syndrome (HGPS) patients, a premature aging disorder. Previously, we and others have characterized the accelerated aging of HGPS cells. Progerin accumulation, caused by a mutation in the lamin A gene (LMNA), distorts the nuclear envelope and triggers multiple cellular alterations associated with aging, including DNA damage, transcriptional dysregulation, and the senescence associated secretory phenotype (SASP) ^54^. Using ECs and VSMCs differentiated from induced pluripotent stem cells derived from HGPS skin fibroblasts, we observed progressive cellular loss and impaired luminal surface integrity in the vascular avatar at day 15, day 30, day 45 and day 90 (Figure 7A, B). VE-cadherin staining revealed intercellular gaps in the HGPS avatar at days 15 and 60 (Figure 7C, D). In addition, nitric oxide production was reduced in the HGPS avatar at day 15 and day 30 (Figure 7E, F), indicating impaired endothelial function. We next detected molecular hallmarks of vascular aging in the HGPS avatar. Both mRNA expression and soluble protein levels of the inflammatory adhesion molecules ICAM1 and VCAM1 were increased in the HGPS avatar compared with the healthy vascular avatar (Figure 7G, H). Furthermore, the expression of SASP associated genes including IL-1b, IL-6, IL-8, and MCP-1 were progressively elevated in the HGPS vascular avatar from day 3 to day 30 (Figure 7I). Immunofluorescence staining of γH2AX and β-gal showed increased proportions of DNA damage–positive and senescent cells respectively in the HGPS avatar compared with the healthy avatar (Figure 7J-M). Of note, these differences became more pronounced at later time points.

**Figure 7.**
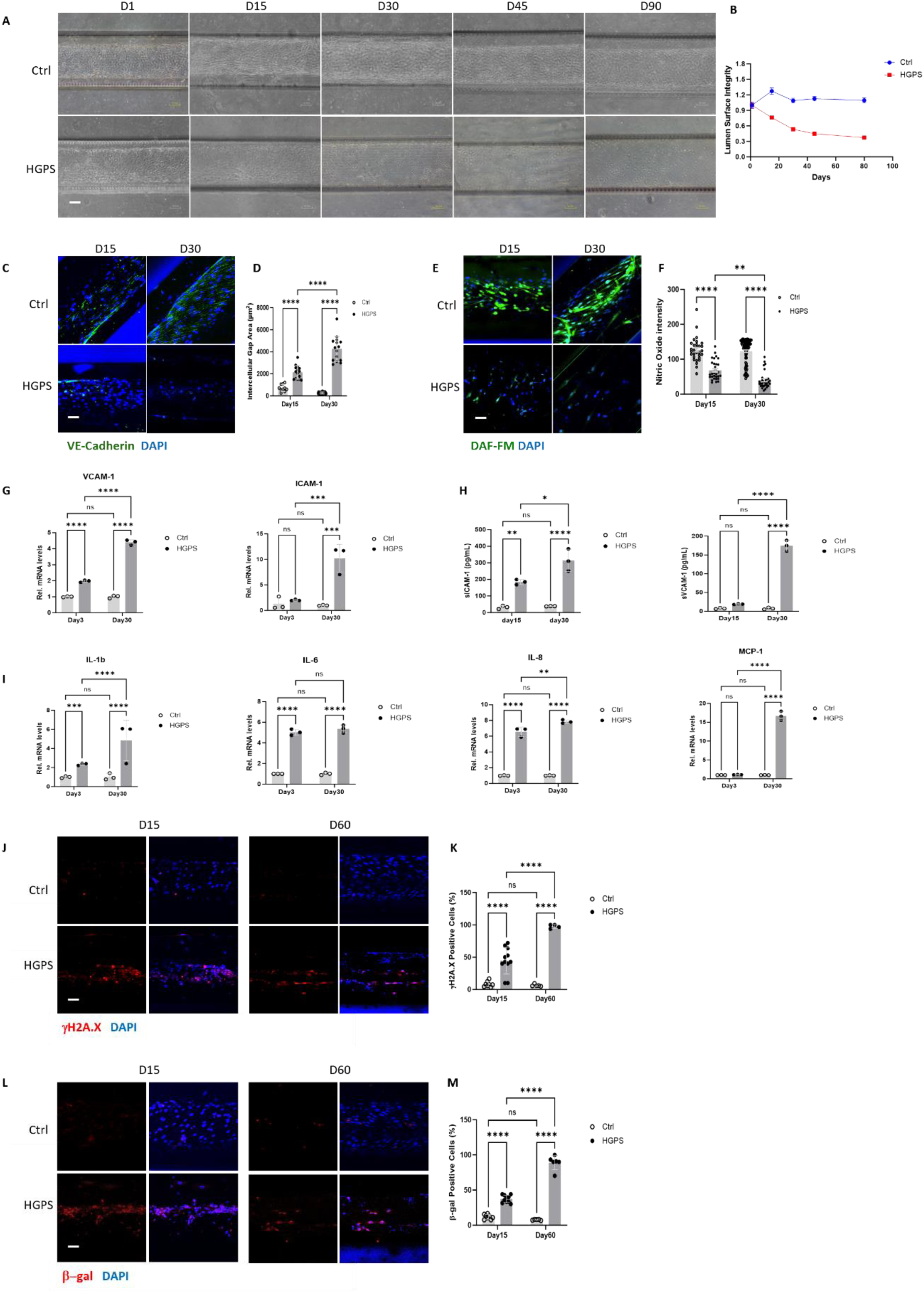
Modeling accelerated vascular aging using the long-term vascular avatar. **A**, Representative images of co-culture of ECs and VSMCs from non-HGPS (Ctrl) or HGPS at the indicated time points (scale bar: 100 µm). **B**, Quantification of **(A)** shows progressive disruption of lumen surface integrity throughout the 90-day co-culture of HGPS ECs and VSMCs, compared to co-culture of non-HGPS (n=6). **C**, Immunofluorescence staining shows loss of the endothelial marker VE-Cadherin in co-culture of ECs and VSMCs from HGPS, compared with co-culture of non-HGPS (scale bar: 50 µm). **D**, Quantification of **(c)** indicates increasing intercellular gap area at day 15 and day 30 in co-culture of HGPS ECs and VSMCs, compared with non-HGPS. Each dot represents one intercellular gap area measured from 3 vascular avatars. **E**, Immunofluorescence staining shows reduced nitric oxide levels, measured by DAF-FM fluorescence, in co-cultures of HGPS ECs and VSMCs at day 15 and day 30 compared with non-HGPS (scale bar: 50 µm). **F**, Quantification of **(E)** shows reduced nitric oxide signal intensity in co-culture of HGPS ECs and VSMCs at day 15 and day 30, compared with non-HGPS. Each dot represents one cell from 3 vascular avatars. **G,** qPCR analysis shows increased inflammatory markers VCAM-1 and ICAM-1 in HGPS co-culture at day 15 or day 30 (n=3). **H**, HGPS co-culture shows increased secretion of soluble sICAM-1 and sVCAM-1 in vascular avatars at day 15 or day 30 (n=3). **I**, qPCR analysis shows increased expression of inflammatory cytokines IL-1b, IL-6, IL-8, and MCP-1 in HGPS co-culture at day 15 and day 30 (n=3). **J**, Representative images of γH2AX staining in vascular avatar at day 15 and day 60 (scale bar: 50 µm). **K**, Quantification of **(J)** shows an increased percentage of γH2AX positive cells in HGPS co-culture. Each dot represents one field from 3 vascular avatars. **L**, Representative images of β-gal staining in vascular avatar at day 15 and day 60 (scale bar: 50 µm). **M**, Quantification of **(L)** shows an increased percentage of β-gal positive cells in HGPS co-culture. Each dot represents one field from 3 vascular avatars. Data between two groups were analyzed using Student’s t-test and comparison among multiple groups were analyzed by two-way ANOVA. Statistically significance defined as P<0.05(*), P<0.01(**), P<0.001(***), and P<0.0001(****).

To further characterize cellular heterogeneity within the vascular avatar, we performed scRNA-seq of both HGPS and non-HGPS avatar at day 30. Based on the marker gene expression, we obtained 13 distinct EC and VSMC sub-clusters (Figure S8A-C). For further interpretation, these subpopulations were grouped into four major categories (Figure S8D,): (1) functional ECs, including angiogenic ECs and mature ECs; (2) dysfunctional ECs, including stressed ECs, activated ECs, EndMT EC1, and EndMT EC2; (3) functional VSMCs, represented by contractile VSMCs; and (4) dysfunctional VSMCs, including synthetic VSMC1, synthetic VSMC2, ECM VSMCs, epithelial VSMCs, aPMC, and cycling cells. Heatmap analysis of representative marker genes confirmed that functional ECs expressed higher levels of endothelial identity genes such as VEGFA, KDR, PECAM1, CDH5, ENG, vWF and CLDN5, whereas dysfunctional ECs exhibited elevated expression of inflammatory molecules such as ICAM-1, CCL2 and CXCL8. Similarly, functional VSMCs were characterized by high expression of contractile markers such as TAGLN and ACTA2, while dysfunctional VSMCs express markers associated with hyperplasia and extracellular matrix synthesis, including COL3A1, PDGFRB (Figure S8A; Figure 8A).

**Figure 8.**
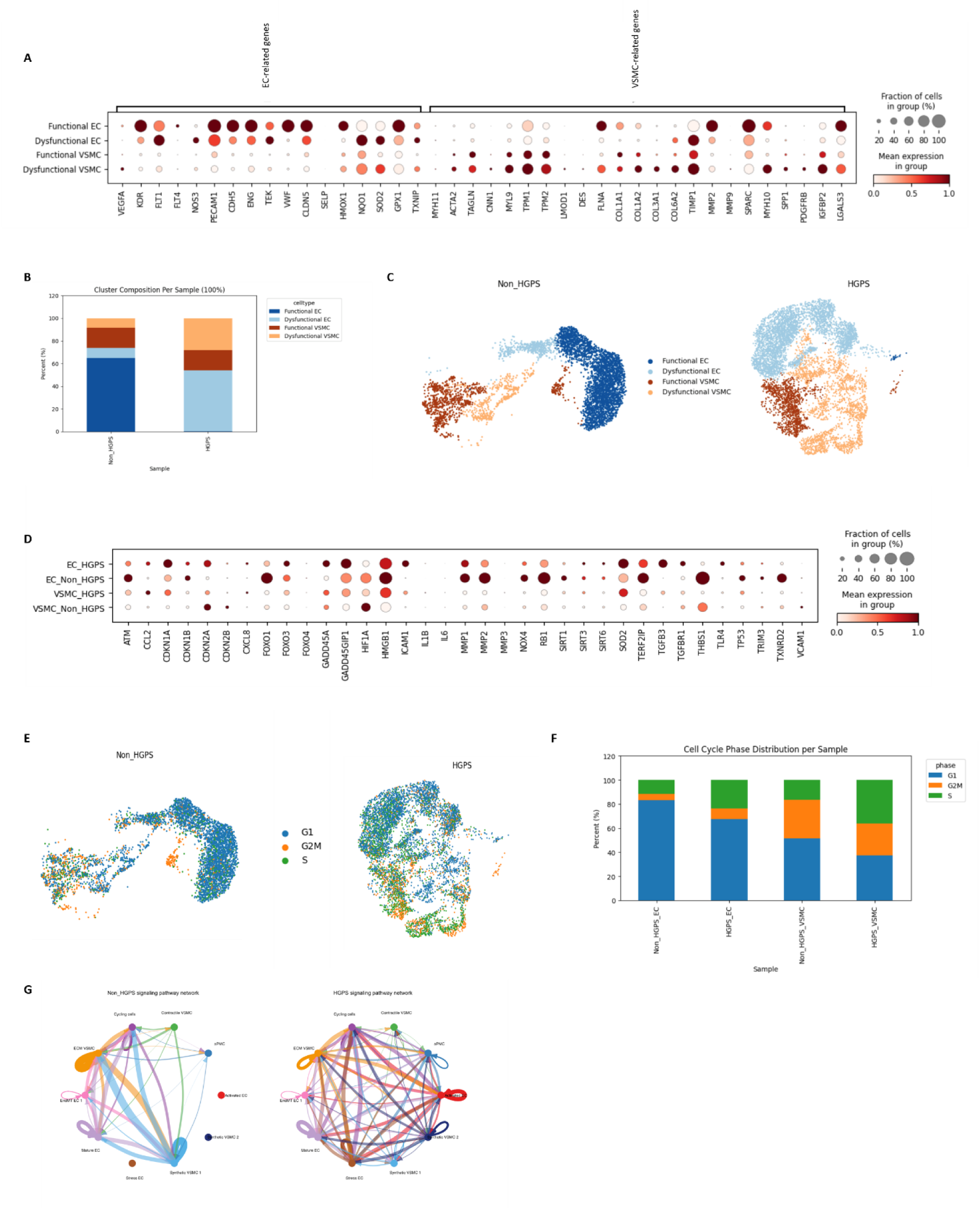
Single cell analysis of accelerated aging in the vascular avatar. **A**, Representative marker genes identifying functional ECs, dysfunctional ECs, functional VSMCs and dysfunctional VSMCs. **B**, Percentages of functional ECs, dysfunctional ECs, functional VSMCs and dysfunctional VSMCs from non-HGPS and HGPS at day 30. **C**, UMAP plots showing functional ECs, dysfunctional ECs, functional VSMCs and dysfunctional VSMCs detected in non-HGPS and HGPS vascular avatars at day 30. **D**, UMAP plots based on cell cycle scores for non-HGPS and HGPS are shown. **E**, Percentages of ECs and VSMCs in different cell cycle phases from non-HGPS and HGPS at day 30. **F**, Representative expression of senescence marker genes in ECs and VSMCs from non-HGPS and HGPS at day 30. **G**, Predicted cell-cell communication between ECs and VSMCs sub-populations.

Notably, functional ECs accounted for over 60% of ECs in the non-HGPS avatar, whereas dysfunctional ECs were over-represented in the HGPS avatar. Consistent with these observations, heatmap analysis of senescence-associated genes revealed increased expression of CDKN1A (p16), CDKN2A (p21), CXCL8, ICAM-1 and TLR4 in ECs from the HGPS avatar compared to non-HGPS avatar (Figure 8D). Dysfunctional VSMCs were greatly increased in the HGPS avatar (Figure8B, C). Cell cycle analysis further revealed that a higher proportion of ECs and VSMCs from the HGPS avatar were found in the S phase (Figure S8E; Figure 8E, F), indicating increased DNA replication and cell cycling. Moreover, cell–cell communication analysis indicated enhanced intercellular crosstalk among 13 sub-populations in the HGPS avatar compared with non-HGPS avatar (Figure 8G). Collectively, these findings demonstrate that the long-term vascular avatar recapitulates key features of vascular aging and provides a novel platform for modelling vascular disease.

## Discussion

While vessel-chip technology holds great promise for studying vascular disease and facilitating drug discovery, maintaining normal structure and function of a vascular co-culture over extended periods has remained a major challenge. We have developed a human vascular avatar that can be maintained for over six months while maintaining vascular morphology, function and transcriptional profile. Our approach was to begin by optimizing conditions for the endothelial monolayer. Our rationale was that the endothelium exerts substantial control over vessel homeostasis, generating factors such as nitric oxide that maintain EC identity and viability, and which suppress the expression of inflammatory cytokines, and the contractility and proliferation of the underlying vascular smooth muscle^55,56^. A panel of three endothelial biologists, supported by evidence from the literature, selected a cohort of potential agents to maintain long-term EC cultures, and were guided by the effects of these agents on cellular viability, morphology, and transcriptional signatures.

A major problem with long-term culture of ECs is the loss of cell identity over time. For example, EndoMT commonly occurs during long-term culture, likely driven by inflammation, oxidative stress and replicative senescence. We observed that in standard EC culture conditions, approximately 30% of HAECs exhibited EndoMT, with VE-Cadherin and α-SMA double-positivity at day 40 (Figure 1G, H). This loss of cell identity was accompanied by elevated inflammatory markers IL-1b, IL-8, ICAM-1, and VCAM-1. However, cells exposed to standard EC medium plus the with TGF-β inhibitor (SB 431542, 10 μM), manifested less EndoMT in long-term culture (Figure 1E-H). In addition to inhibiting EndoMT, TGF-β inhibition would be expected to oppose cellular senescence^57–60^. This is because TGF-*β* is known to promote cellular senescence, mediated in part by reducing H4K20me3 via miR-29 ^59^. Indeed, the SB compound did suppress morphological manifestations of cellular senescence (increased cell area and cell circularity). However, medium containing the TGF-β inhibitor did not entirely suppress inflammatory responses (Figure S1C, D,).Similarly, VEGF is a known promoter of EC identity and survival, and enhances EC migration, proliferation and network formation^61,62^. Addition of VEGF to the medium enhanced EC viability and proliferation, and reduced the expression of inflammatory cytokines. However, by day 60 the EC monolayer was overpopulated, cell margins were difficult to identify, and RNAseq suggested that a cell fate transition was underway, with neuronal pathways represented (Figure S1B). Other than low serum conditions, none of the other factors had favorable effects on long-term EC culture. Because treatment with medium containing VEGF or SB compound had complementary effects, it seemed sensible to combine the treatments. We found that combinatorial treatment with VEGF, SB, and low FBS (VSL) preserved EC identity and function during long-term culture, and suppressed senescence and inflammation. This optimized condition supported 3D human EC cultures for over 180 days, maintaining healthy morphology, stable barrier integrity, and a non-inflammatory secretome (Figure 3).

Subsequently, we assessed this optimized medium (VSL) in co-culture. To our dismay, the VSL media was not optimal for co-culture; by day 30 the luminal integrity of the co-culture was compromised by over-proliferation of the VSMCs. Accordingly, we further optimized the VSL medium conditions to prevent VSMC over-proliferation and maintain a quiescent state. Because FGF is a known agonist of VSMC proliferation^63^, we assessed the effect of depleting it from the medium. Because PDGF is a known paracrine agonist of proliferation, we also studied the effect of adding a PDGF inhibitor^47^. Because prostacyclin and its second messenger cAMP are potent suppressors of VSMC proliferation, we also studied the effect of the stable analogue, 8Br-cAMP^49^. Finally, we further reduced the serum concentration in the medium. Whereas each of these conditions reduced VSMC proliferation and inflammatory signaling, the combination of these agents was most effective, as assessed by the long-term maintenance of a mature VSMC culture as assessed by expression of mature VSMC markers and inflammatory cytokines (Figure 4). The mVSL+ medium successfully established these conditions.

Human blood vessels are continuously exposed to numerous circulating cardiovascular disease (CVD) risk factors that contribute to pathology: metabolic factors (glucose, homocysteine, lipids), hormones (insulin, endothelin-1, angiotensin II), and inflammatory cytokines. For example, high blood glucose levels impair endothelial vasodilator function^64^ and induce DNA damage, inflammation, mitochondrial dysfunction and ROS accumulation^39,65^. The exposure to hypercholesterolemia impairs endothelial functions, elevates blood pressure and accelerates atherosclerosis^66,67^. Similarly, elevated blood levels of Angiotensin II triggers hypertension, potentially causing heart failure in part by inducing EC senescence^39^ ^68^ ^69^. Current in vitro models do not facilitate the study of the long-term effects of risk factors on the endothelium and vascular smooth muscle. In contrast, our long-term vascular avatar enables perturbation studies with real-time monitoring of morphology, luminal integrity, and the vascular secretome, and—using iPSC-derived vascular cells—supports patient-specific modeling toward personalized medicine, including vascular stressors such as cosmic radiation and microgravity.

Although a few studies have incorporated HGPS vascular cells into microphysiological platforms, these cultures were maintained for less than one week ^70,71^, which limits the ability to study chronic alterations such as senescence. In our HGPS avatar, differences in determinants of structural junction integrity (Reflected by VE-Cadherin immunofluorescence), nitric oxide production, inflammatory signaling, DNA damage, and senescence/SASP markers progressed over time, and became pronounced by day 30 (Figure 7). Single-cell RNA sequencing at day 30 indicated that ∼75% of HGPS vascular cells exhibited a dysfunctional phenotype (Figure 8). Notably, ∼40% of VSMCs remained functional. This observation contrasts with the prevailing view that VSMC pathobiology drives HGPS vascular disease ^72–74^.

One limitation of our study is that we did not include physiological hemodynamic factors, with our co-culture in a low shear stress state. This may explain the phenotypic shift in the ECs of the vascular avatar over time, from an arterial phenotype to a venous/lymphatic EC subtype as assessed by scRNAseq. In the future we intend to introduce physiological shear stress and cyclic strain to mimic in vivo hemodynamics.

## Concluding remarks

We have successfully engineered and maintained a human vascular avatar for over 180 days, using an experimentally derived medium containing VEGF, 8Br-cAMP, the viability enhancer HMH 1015, and inhibitors of TGF-β (SB431542), and PDGF (AG1295). These factors were formulated in basal endothelial medium lacking FGF and supplemented with low concentration of fetal bovine serum (FBS, 1%), together referred to as mVSL+ supplement. Our data indicates that this optimized medium preserves cellular identity, reduces apoptosis and senescence, and prevents excessive VSMC proliferation. Furthermore, we used the avatar to model vascular aging using iPSC-derived vascular cells from patients with HGPS. The HGPS vascular avatar exhibited gradual loss of luminal surface integrity, increased intercellular gap formation, and reduced EC function by day 30. This disease phenotype was accompanied by progressively increased expression of senescence-associated markers and inflammatory mediators. These alterations recapitulate hallmark features of progeria, characterized by endothelial and VSMC dysfunction leading to progressive vascular aging. Collectively, this study demonstrates that the vascular avatar provides a robust and physiologically relevant platform for modeling disease-associated vascular aging and for evaluating therapeutic strategies targeting chronic vascular disorders.

## Acknowledgments

We acknowledge the Advanced Cellular and Tissue Microscopy (ACTM) Core Facility of the Houston Methodist Research Institute for assistance in cellular imaging. We acknowledge HICOMP for their collaboration with the fabrication of Vessel-Chips. We thank Kiran Kuriakose for his thoughtful editing of the manuscript.

## Author’s contributions

JPC, AM, AJ and WQ conceived the study and designed the experiments. WQ and KT performed the experiments and generated the figures. GW, AM, AT, IK, YX and WQ analyzed the bioinformatics data. TKCN and AM generated iPSC-ECs. KTC, VVS and KWB conducted scRNAseq. AK, RP, and BMAD helped analyze data and fabricated devices. WQ and KT generated a draft of the manuscript which was reviewed and revised by all authors. JPC, AJ and GW supervised the personnel and funded the study. JPC provided final review and approval of the manuscript.

## Methods and Materials

### 2D cell culture

Plates or dishes were coated with 0.2% gelatin (Cat# 0423). HAECs or iPSC-ECs were used for cell studies. HAECs were purchased from ATCC (Cat# PCS-100-011). iPSC-ECs were generated using established methods in our laboratory ^75^. Standard EC media (Lonza, EGM-2MV, Cat# CC-4147) was used as a control for modifications that were made to that media to enhance cellular longevity. Cells were incubated at 37°C and 5% CO_2_ (Thermo Scientific) and culture media was changed every other day.

### Fabrication of Vessel-Chips for EC monolayer growth

Vessel-Chips were placed on a sterile plate (15-cm plate) and then placed into an oxygen plasma device to sterilize and activate the surface of the chips. The Vessel-Chips were coated by applying 10-20µL mixture of collagen and fibronectin with a final concentration of 1mg/mL collagen (Cat# 354249) and 50 µg/mL fibronectin (Cat# CB-40008A) and incubated at 37°C for 1 hr. ECs (20ul from a mixture of 10 × 10^6^ cell/mL in complete medium) were seeded in Vessel-Chips after washing with 1X PBS. The Vessel-Chips with cells were placed on a rotator and incubated at 37°C for 3 hr. The chips were washed with 200 µL complete medium to remove non-attached cells and incubated at 37°C and 5% CO_2_ (Thermo Scientific). Media was changed daily.

### Fabrication of Vessel Chips for Co-culture using Gravitational Lumen Patterning (GLP)

GLP Vessel-Chips were fabricated as previously described ^14,15^. A coating solution was prepared by diluting rat tail type I collagen (10 mg/mL, Ibidi) to a final concentration of 5 mg/mL and adjusting the pH to 7.2–7.4 using sodium bicarbonate (7.5%, Gibco), sodium hydroxide (Thermo Fisher), and HEPES (1 M, Gibco). VSMCs were suspended in culture medium to a final concentration of 5 × 10⁶ cells/mL and gently mixed with the collagen solution to ensure uniform distribution. The pH was verified by placing a drop of the mixture onto pH paper.

Prior to lumen patterning, 200-µL pipette tips (P200) were bent to approximately 135° at ∼7 mm from the distal end. A straight P200 tip was used to aspirate ≤20 µL of the collagen–VSMCs mixture and inject it into the outlet port of the microfluidic channel until the collagen levels were approximately equal at both the inlet and outlet. A second bent P200 tip containing 20 µL of 1× PBS was inserted into the inlet port. The devices were then held vertically with the outlet ports facing downward and immediately transferred to a humidified incubator (37 °C, 5% CO₂).

After 7 minutes of incubation, the devices were returned to the biosafety cabinet. Outlet tips were removed by gently twisting while keeping the chips upright. The chips were then placed horizontally, the inlet tips removed, and 200 µL of culture medium added to each chip. Chips were incubated at 37 °C with 5% CO₂. Twenty-four hours after VSMCs loading, ECs were seeded onto the lumen surface as described in the section on endothelial monolayer formation. Culture medium was replaced daily by aspirating the old medium and adding fresh medium through the same inlet ports.

### Maintenance of human induced pluripotent stem cells

Human iPSC lines (HGFDFN168 iPSC1P, HGADFN167 iPSC 1Q) provided by the Progeria Research Foundation Cell and Tissue Bank, were cultured on Matrigel-coated plates (BD Biosciences; Corning plates) in mTeSR Plus medium (STEMCELL Technologies) following the manufacturer’s protocol and our previous work^38^. The iPSCs were passaged upon 70% confluency and maintained in humidified incubators at 37°C and 5% CO2 (Thermo Scientific).

### Generation of endothelial cells from induced pluripotent stem cell (iPSC)

Endothelial cells were differentiated from iPSCs using our well-established protocol as previously described^38,43^. Briefly, iPSCs at ∼60% confluence were cultured in DMEM/F12 medium supplemented with the Wnt agonist CHIR99021 (5 μM, Selleck), bone morphogenetic protein 4 (BMP4, 25 ng/mL, Peprotech), B27 supplement (Thermo Fisher Scientific), and N2 supplement (Thermo Fisher Scientific). On day 3, cells were detached using Accutase, re-seeded onto Matrigel-coated plates, and cultured in StemPro medium supplemented with forskolin (5 μM, LC Laboratories), VEGF (5 ng/mL, Peprotech), and polyvinyl alcohol (2 mg/mL, Sigma).

On day 7, the medium was replaced with complete endothelial growth medium (EGM-2MV, Lonza) supplemented with 100 ng/mL VEGF. On day 11 or 12, ECs were isolated using CD31 microbeads. Purified iPSC-derived ECs were either maintained in culture or used for downstream experiments after day 12.

### β-Galactosidase staining

β-Galactosidase staining was performed according to the manufacturer’s protocol (Cell Signaling Technology, Cat# 9860). Briefly, cell culture medium was removed, and cells were rinsed once with 1x PBS. Cells were then incubated with 1x fixative solution for 15 min at room temperature. After fixation, cells were rinsed twice with 1x PBS and incubated with 1x β-gal staining solution at 37°C overnight in a dry, CO₂-free incubator. Images were acquired using an inverted microscope (Nikon).

### Measurement of Nitric Oxide Production

At day 30 after treatment with different circRNA hTERT, vascular avatars were collected for staining with DAF-FM fluorescent dye (Cat# D-23841, Invitrogen) (Mojiri et al. 2021). Briefly, vascular avatars were incubated with 5 μM DAF-FM dye in standard EC media for 30 min at room temperature. The dye was washed away after incubation, and standard EC media was added to allow the complete de-esterification of AM moieties and nitric oxide-dependent fluorescence development. Images were acquired using a 10X fluorescence microscope (EVOS M5000) and analyzed with ImageJ. Signal intensity was quantified as the mean fluorescent intensity per cell.

### Bio-Rad ELISA analysis

The concentrations of inflammatory cytokines in EC-conditioned media were measured using the Bio-Plex Pro™ Human Cytokine assay (Bio-Rad Laboratories, Hercules, CA, USA) following the manufacturer’s instructions. EC media was collected at designated time points and either stored at -30°C or used immediately for analysis. Two inflammatory markers (soluble ICAM-1 and VCAM-1) or 48 cytokines were detected using the corresponding primary antibodies. Beads were washed three times with 1x wash buffer and incubated for 1h with fluorescently labeled secondary antibodies of distinct emission profiles. After a final wash, assay buffer was added, and the plates were read using the Luminex 200 System.

### Real-Time PCR analysis

Cell pellets were collected, and total RNA was extracted using a RNeasy Mini Kit (QIAGEN) according to the manufacturer’s protocol. cDNA was synthesized from the 100 ng of RNA per reaction using the 5x iScript Reverse Transcriptase Supermix (Bio-Rad Laboratories). Each qPCR reaction (10 µL total volume) contained 5 µL Power SYBR Green Master Mix (Applied Biosystems, Thermo Fisher Scientific), 0.5 µL of forward and reverse primers, 1 µL of cDNA, and 3.5 µL of nuclease-free water. Quantitative real-time PCR was performed using the QuantStudio 12K Flex system (Applied Biosystems, Life Technologies). Gene expression levels were calculated relative to control samples using the ΔΔCt method ^76^. Primer sequences used in this project are listed in Table 1.

### Immunofluorescence staining

Immunofluorescence staining for VE-Cadherin staining (Cat# sc-9989), α-SMA staining (Cat# ab5694), Calponin-1 staining (Cat# MA5-11620), β-gal staining (Cat# 66586-1-Ig), and γH2A.X staining (Cat# JBW301) was performed with modifications from our previously published protocol ^38^. Cells were cultured on chamber slides and fixed with 4% paraformaldehyde in PBS for 10 min at room temperature. Permeabilization and blocking were carried out simultaneously by incubating the cells in 0.15% Triton X-100, 0.1% Tween-20, and 3% BSA in PBS for 60 min.

Samples were incubated with primary antibodies diluted in 3% BSA in PBS overnight at 4°C in a humidified chamber. The next day, cells were incubated with secondary antibodies diluted in 3% BSA for 2h at room temperature in the dark. After three washes with 1x PBS, cells were mounted using DAPI (50 µL), then sealed with a coverslip and nail polish. Images were obtained using a FV3000 confocal microscope at the ACTM Core.

### Measurement of lumen surface integrity

Lumen integrity was measured using the Coherence parameter from the Orientation J plugin. This analysis quantifies EC coverage at the lumen midplane by measuring the continuity of the dark shadow produced by a confluent endothelial monolayer under phase-contrast microscopy. Because the lumen has straight edges, aligned ECs form a continuous dark line along the channel. Loss of endothelial integrity (due to apoptosis or cell detachment) reduces this continuity. Thus, the Coherence metric serves as a proxy for lumen integrity by quantifying how well cells maintain alignment along the patterned channel edges.

### Bulk RNA-seq data analysis

Gene expression tables generated by Cuffdiff were merged into a single expression matrix using matching gene identifiers and transformed using log₁₀ (FPKM + 1). Genes detected in fewer than three samples were removed. Batch correction across sample groups was applied only for dimensionality reduction and visualization. Principal component analysis (PCA) was performed on the batch-corrected matrix, followed by nonlinear embedding of the PCA coordinates to visualize transcriptome-wide relationships. Differentially expressed genes (DEGs) were obtained from the uncorrected Cuffdiff outputs (q < 0.05, |log₂FC| ≥ 0.5) and analyzed using a web-based GO functional annotation tool with FDR-adjusted significance. Curated gene panels associated with endothelial identity, inflammation, and senescence were extracted from the uncorrected matrix, and heatmaps were generated from row-normalized log₁₀(FPKM + 1) values with clustering of both genes and samples.

### Preparation for Single Cell Studies

Cells were isolated from Vessel-Chips using TrypLE (Gibco, Cat# 12605010) and incubated at 37°C for 5 mins. The dissociated cells were collected in 1.5ml DNA LoBind Eppendorf tubes then centrifuged at 300xg for 3 mins at room temperature. Pellets were washed twice with wash solution (1xPBS and 0.04% BSA). Cells were resuspended in 100ul wash solution, kept on ice and counted using an automated Cell Counter. Cell numbers are adjusted according to 10x Genomics Single Cell 3’ Gene expression V3.1 protocol (10x Genomics Protocol #CG100058), and the suspension was diluted to 1000 cells/µl. Single cells were captured with the 10x Chromium controller (10x Genomics) according to the manufacturer’s instructions (document CG000315 Rev D) and as described earlier^77^. Cells were loaded onto a 10x Chip and cDNA was generated using the Chromium Next GEM Single Cell 3’ Kit v3.1 (Dual Index). Libraries were sequenced on the NextSeq 2000 Illumina platform to a targeted depth of 50,000 reads per cell.

### Single cell RNA-seq data analysis

Cells from both samples were filtered based on transcript counts (250–20,000 UMIs) and mitochondrial read percentage (<25%). Data were log-normalized and highly variable genes were selected. Batch effects were corrected using Harmony, followed by PCA on the Harmony-corrected embeddings. Clustering was performed using a shared nearest neighbor (SNN) graph, and UMAP visualizations were generated from the first 30 Harmony components. Cluster markers were identified using FindAllMarkers function in Seurat and annotated according to canonical cell type markers.

### Data analysis

GraphPad prism 10.0 was used for all statistical analyses. Data were presented as Mean ± Standard Error of Mean (Mean ± SEM). Comparisons between two groups were performed using Student’s t-test. Comparisons among multiple groups were analyzed using one-way or two-way ANOVA as appropriate.

### qPCR primers

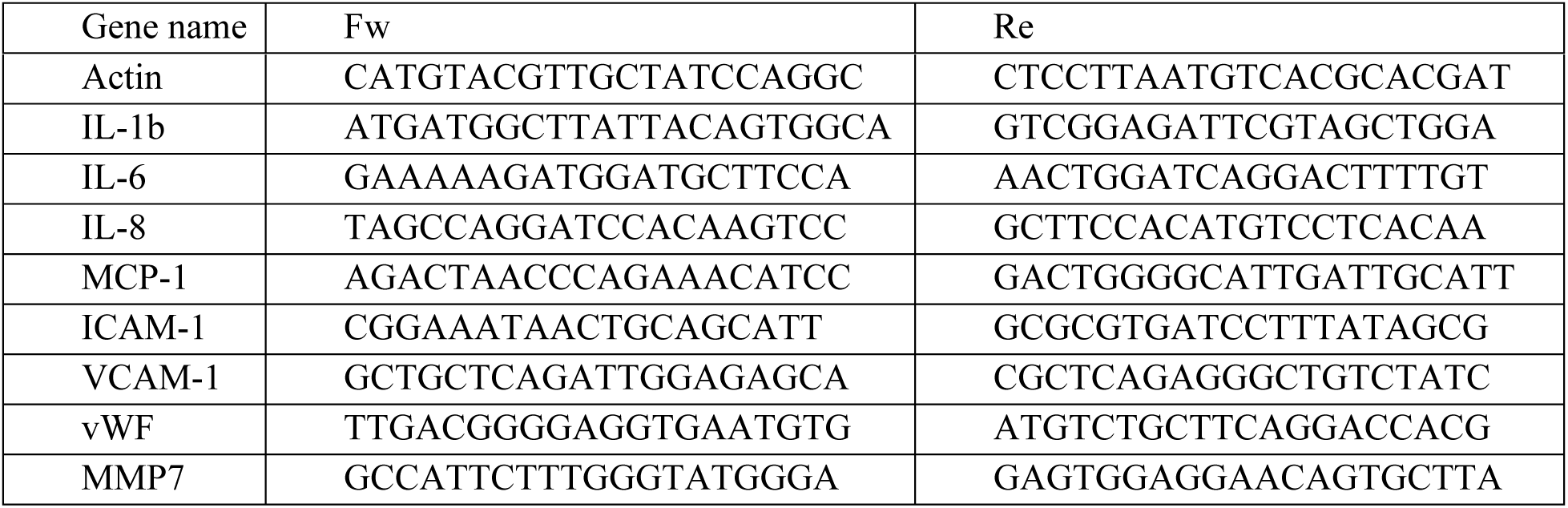

## Abbreviations

GLP: Gravitational Lumen Patterning
ECM: Extracellular matrix
VSMCs: Vascular smooth muscle cells
ECs: Endothelial cells
TGF-β: Transforming growth factor-β
EndoMT: Endothelial-to-mesenchymal transition
PDGF: Platelet-derived growth factor
FGF2: Fibroblast growth factor-2
NO: Nitric oxide
FBS: Fetal bovine serum
HAECs: Human aortic endothelial cells
PCA: Principal Component Analysis
vWF: von Willebrand factor
MMP7: Matrix metalloproteinase-7
iPSC-ECs: Induced pluripotent stem cell-derived ECs
mVSL: modified VSL

---

**Figure S1.**
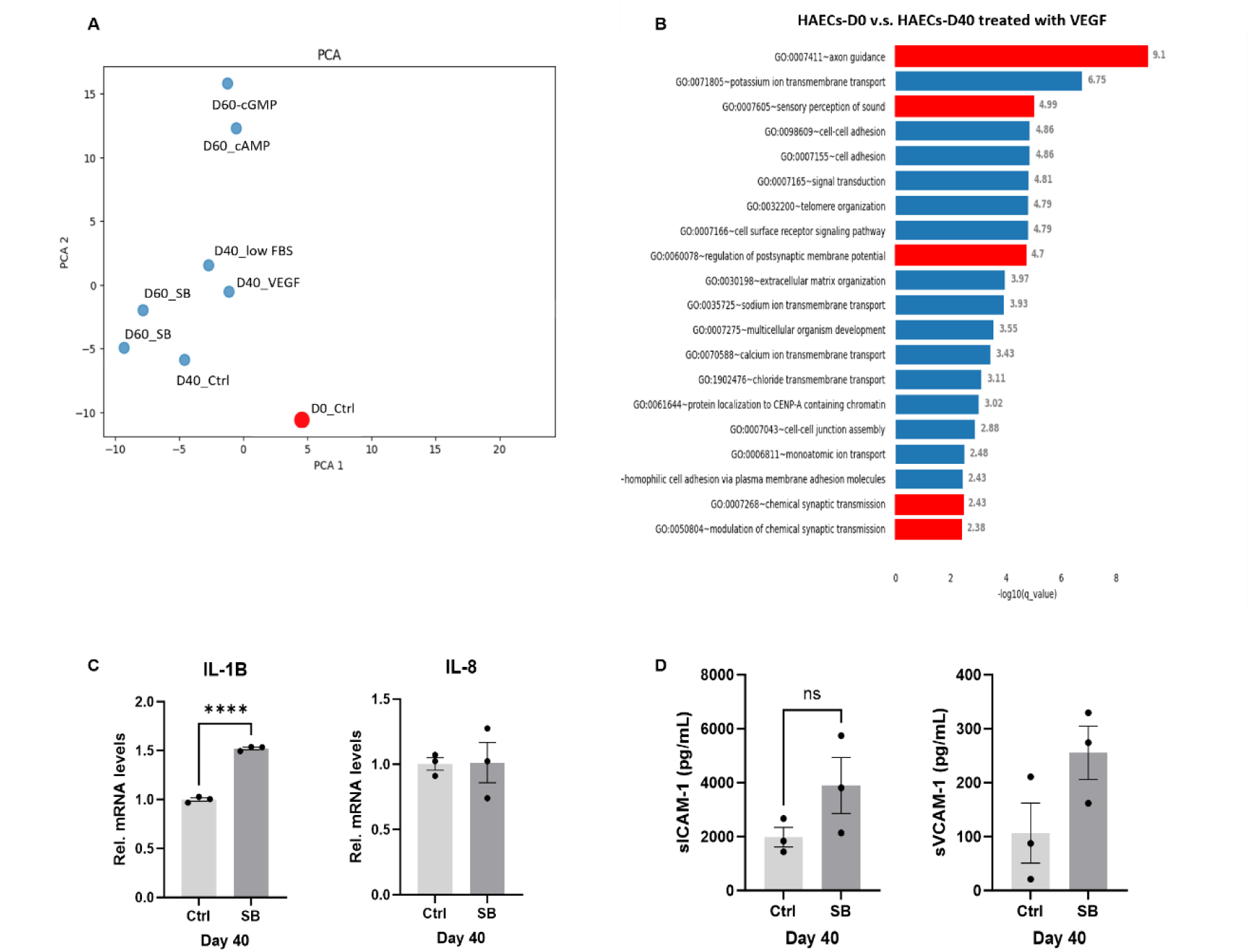
VEGF and SB Differentially Influence EC Longevity and Identity. **A**, UMAP analysis of transcriptional data from EC exposed to standard media (EGM-2MV with 5% FBS) at Day 0 or Day 40 (D0_Ctrl and D40_Ctrl respectively); or standard media plus EC viability factors including TGF b inhibitor SB 431542(SB; 10 µM); vascular endothelial growth factor (VEGF;10 ng/mL); low FBS (3% FBS); 8-Br-cAMP(cAMP;10uM); or 8-Br-cGMP (cGMP, 10uM). **B**, GO analysis for HAECs reveals that additional VEGF (10 ng/mL) in control media (EGM-2MV with 5% FBS) activates neuronal pathways during long-term culture. Neuronal pathways are highlighted in red. **C**, qPCR analysis for HAECs at day 40 reveals that adding SB to the EGM-2MV with 5% FBS elevates inflammatory marker IL-1b (n=3). Each dot represents one technical repeat. Data between two groups are analyzed by Student’s t-test. Results are considered statistically significant with P<0.0001. **D**, ELISA assay for HAECs at day 40 reveals that SB tends to elevate the expression of sICAM-1 and sVCAM-1 in culture media (n=3). Ctrl = standard EC medium. SB = TGF-b inhibitor (SB 431542, 10 µM) added to standard EC medium. Each dot represents one technical repeat.

**Figure S2.**
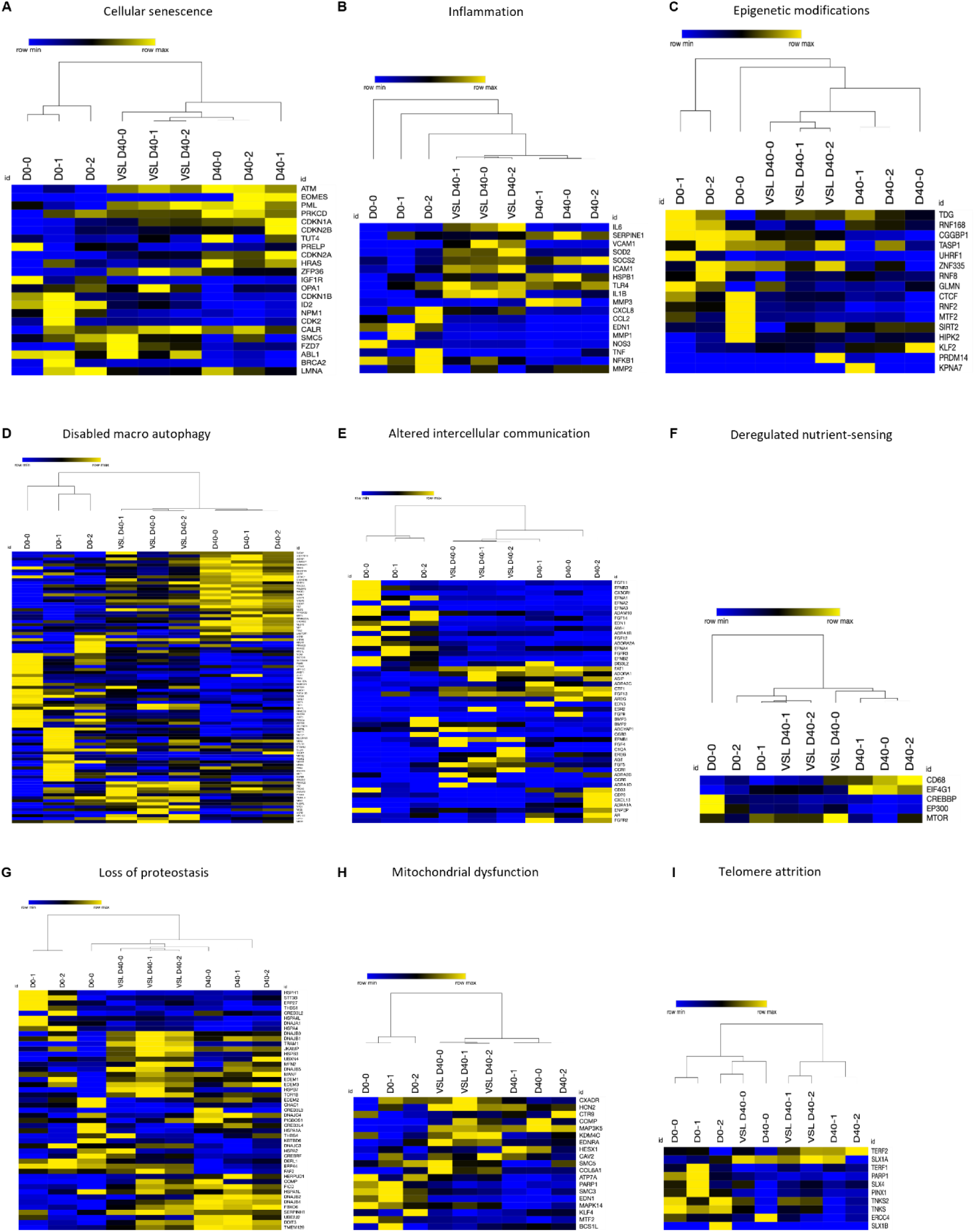
VSL Reverses Hallmarks of Senescence as Revealed by Bulk RNA-seq. Heatmap analysis reveals that VSL reduces expression of genes involved aging processes during long-term culture, including **A**, cellular senescence, **B**, inflammation, **C**, epigenetic modifications, **D**, disabled macro autophagy, **E**, altered intercellular communication, **F**, deregulated nutrient-sensing, **G**, loss of proteostasis, **H**, mitochondrial dysfunction, and **I**, telomere attrition. D0-0, D0-1, D0-2: iPSC-ECs at day 0 in triplicate. VSL-D40-0, VSL-D40-1, VSL-D40-2: iPSC-ECs treated with VSL for 40 days in triplicate. D40-0, D40-1, D40-2: iPSC-ECs treated with standard EC medium for 40 days in triplicate.

**Figure S3.**
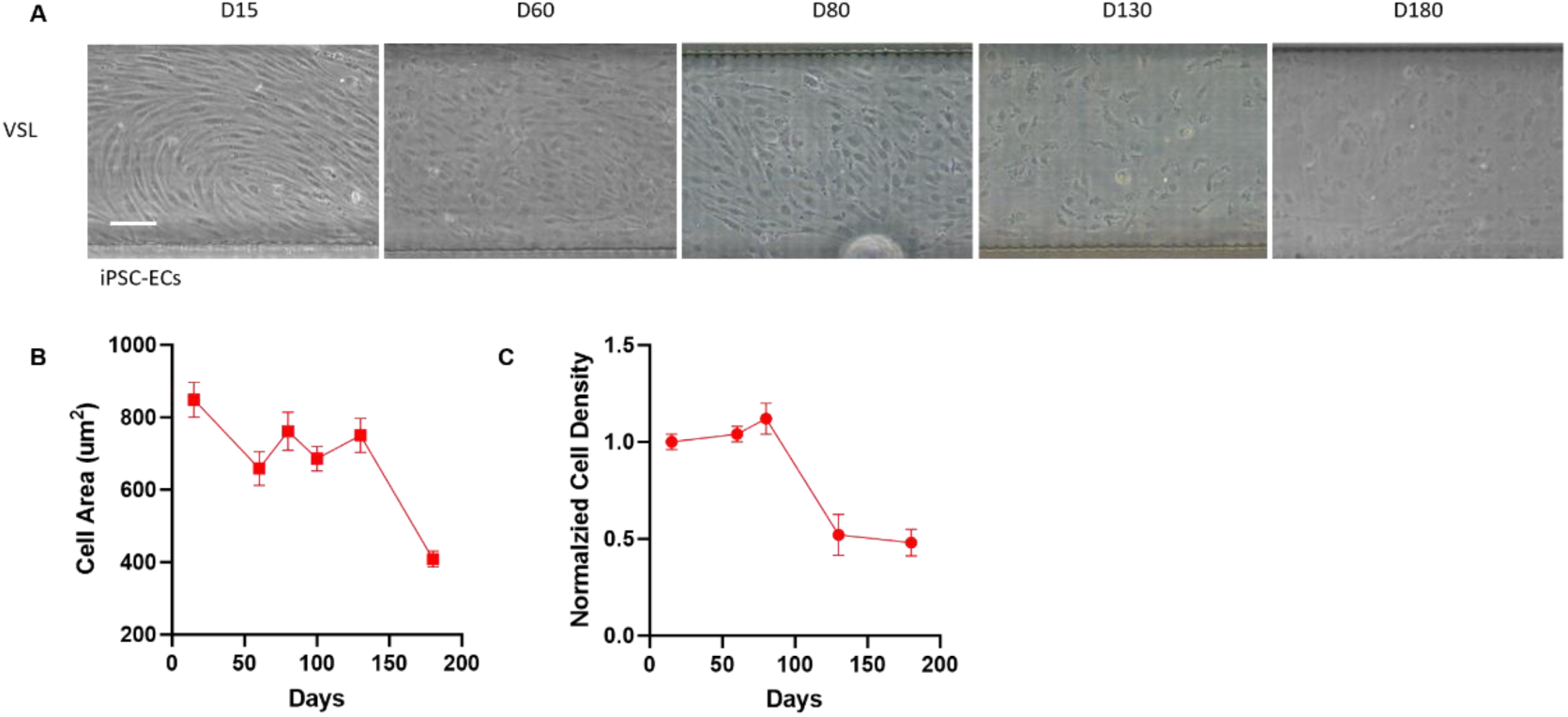
Human iPSC-derived EC monolayer in vessel-chip and morphological characteristics. **A**, VSL sustains EC monolayer derived from iPSC for over 180 days (scale bar: 50 µm). **B**, Quantification of cell area from (**A**) reveals that an increase in cell area (consistent with senescence) is not observed throughout the 180-day culture in VSL media. **C**, However, there is a reduction in cell density.

**Figure S4.**
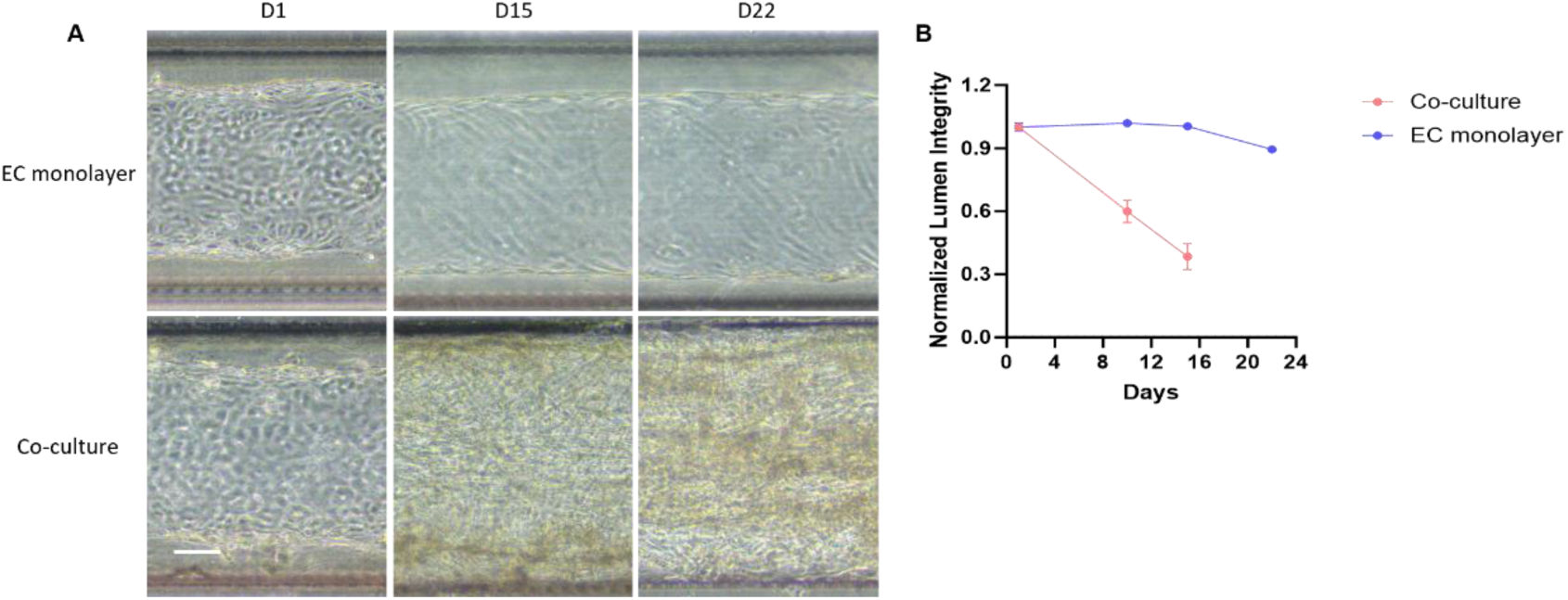
Human vascular avatar co-culture of ECs and VSMC growth in VSL. **A**, Representative images of EC monolayer and VSMC-EC co-culture maintained by VSL in the vascular avatar shows that VSL fails to maintain a co-culture of ECs and VSMCs beyond 30 days (scale bar: 40 µm). **B**, Quantification from (**A**) shows that lumen surface integrity declines in co-culture maintained by VSL within 22 days.

**Figure S5.**
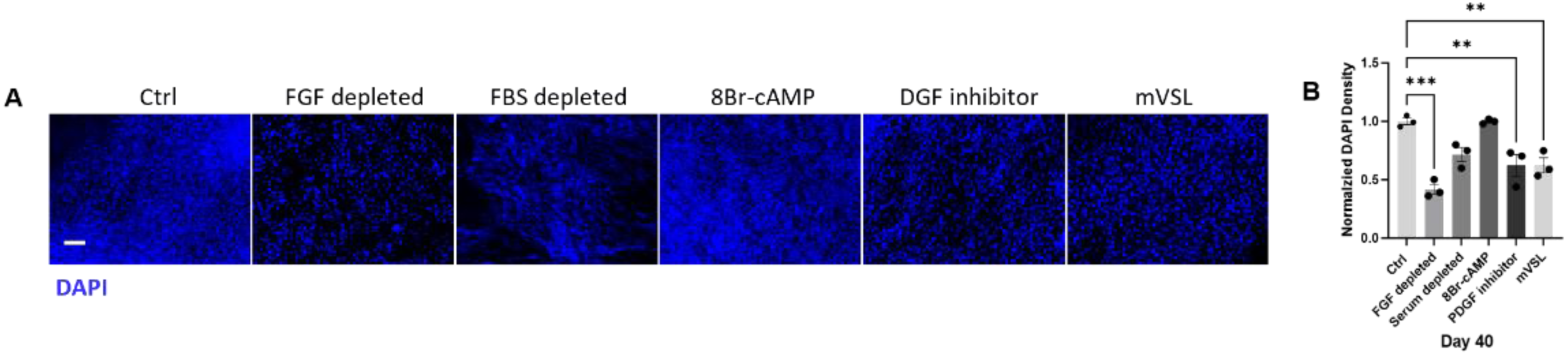
Screening of longevity factors for VSMCs. **A**, Morphometric measurement of iPSC-VSMCs at day 40 reveals that VSMCs treated with mVSL maintain a normal cell profile compared to other groups (scale bar:100 µm). Ctrl = VSL, FGF depleted= VSL with FGF depletion; FBS depleted = VSL with FBS depletion; 8Br-cAMP = 2 µM 8Br-cAMP added to VSL; PDGF inhibitor = PDGF inhibitor (AG 1295, 10µM) added to VSL; mVSL (VSL with PDGF inhibitor (AG1295 10 μM), 8Br-cAMP (2 μM), low serum (1% FBS) with depletion of FGF). (scale bar:100 µm). **B**, Immunofluorescence staining with DAPI shows cell density of each group (scale bar:100 µm). **C**, Quantification of **(B**) by DAPI density reveals that the mVSL reduces the cell density compared to the control group (VSL).

**Figure S6.**
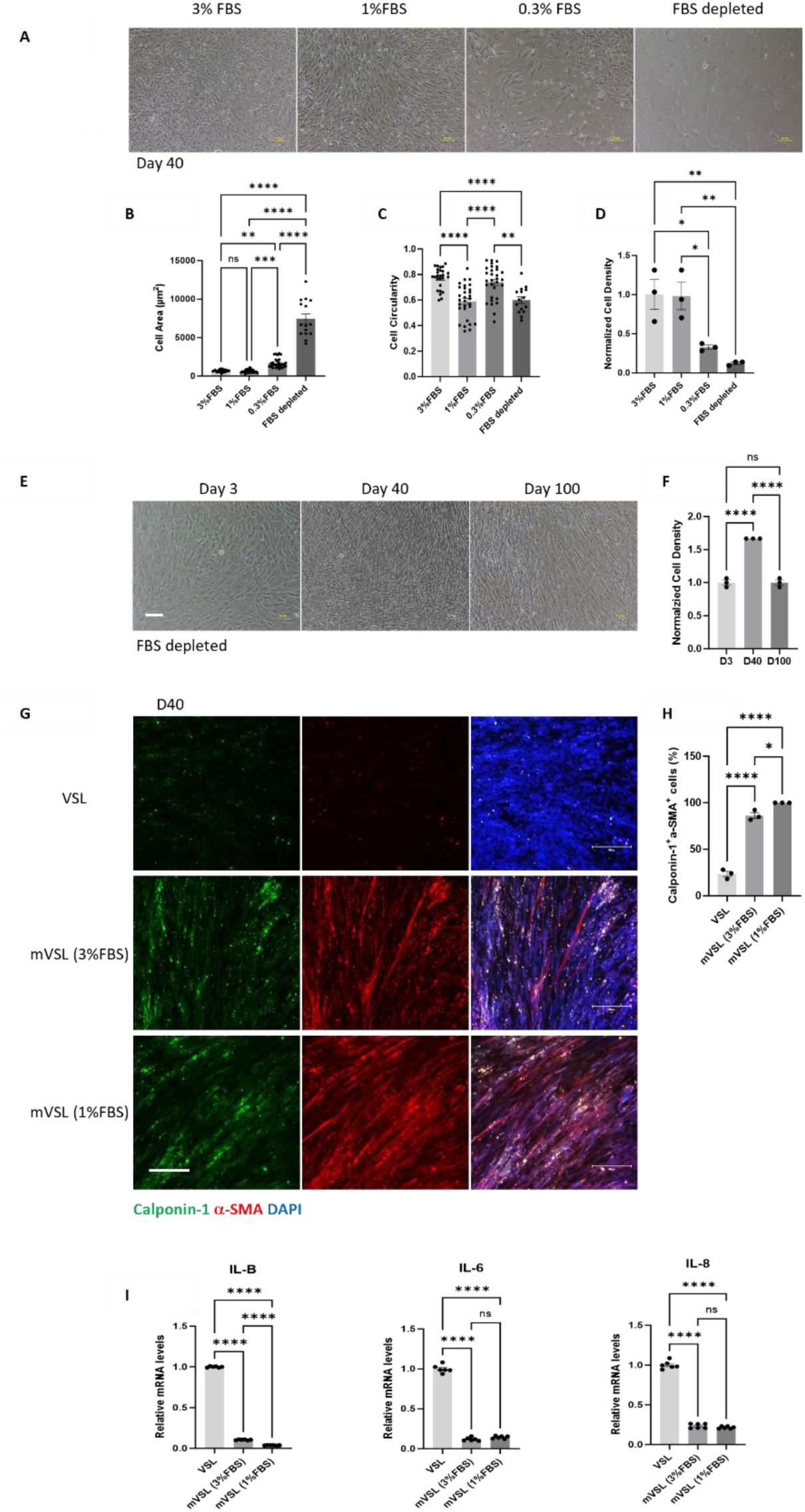
Impact of Reduced Serum Conditions on ECs and VSMCs. **A,** Morphometric measurement shows that reduction of FBS from 3% to 1% or 0.3% in standard EC medium (Lonza, EGM-2MV, 5% FBS) improves EC morphology. However, removal of FBS fails to sustain the viability of ECs for over 40 days. 3% FBS = only 3% FBS added to EC medium; 1% FBS = only 1% FBS added to EC medium; 0.3% FBS = only 0.3% FBS added to EC medium. (scale bar:100 µm). **B**, Quantification of (**A**) shows that removal of FBS increases cell area of HAECs at day 40. **C**, Quantification of (**A)** by cell circularity is shown. **D**, Quantification of (**A**) shows that removal of FBS reduces the cell density of HAECs at day 40. **E**, Morphometric measurement shows that removal of FBS maintains the culture of VSMCs for over 100 days (scale bar:100 µm). **F**, Quantification of cell density from **(E)**. **G**, Immunofluorescence staining shows VSMCs that are Calponin-1 or/and α-SMA positive. mVSL (3% FBS) = only 3% FBS added to mVSL; mVSL (1% FBS) = only 1% FBS added to mVSL. (scale bar: 300 µm). **H**, Quantification of (**G**) reveals that further reduction of FBS to 1% from 3% in mVSL preserves VSMC contractile markers. **I**, qPCR analysis for VSMCs reveals that further reduction of FBS decreased inflammation marker IL-1b.

**Figure S7.**
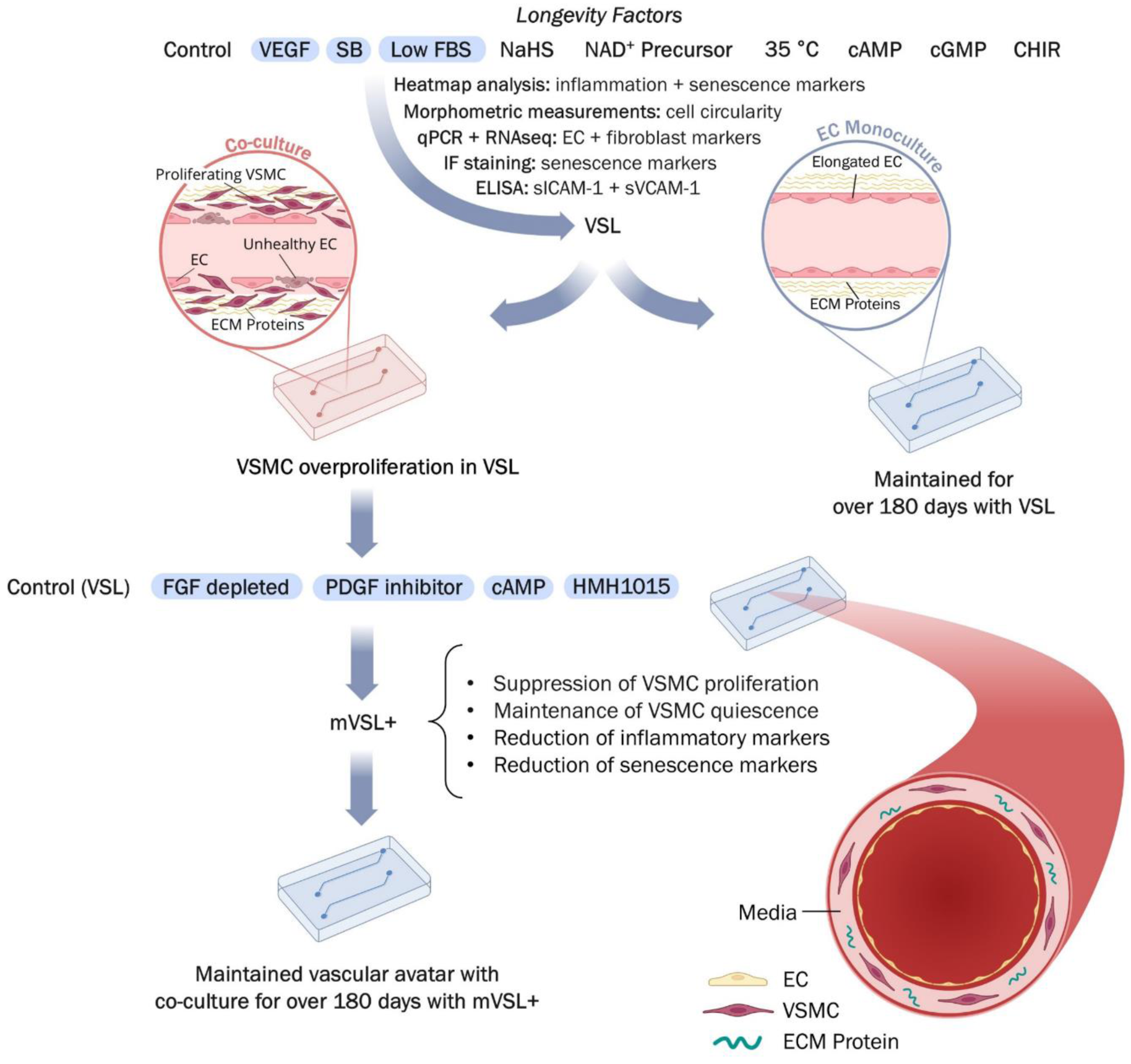
Proposed model for generation of the long-term vascular avatar. A working model shows how this study has been conducted. In brief, we derived three viability factors for long-term culture of ECs: VEGF, SB, and low serum. Combinatorial treatment with these three conditions identified VSL as an optimal culture media for long-term maintenance of an EC monolayer. However, VSL failed to maintain the co-culture for over 30 days, with loss of luminal integrity due to VSMC proliferation. By adding anti-proliferative factors, we were able to generate an optimal media (mVSL+) for the co-culture. Finally, by applying mVSL+ to vessel-chips, we successfully maintained a vascular avatar for over 180 days.

**Figure S8.**
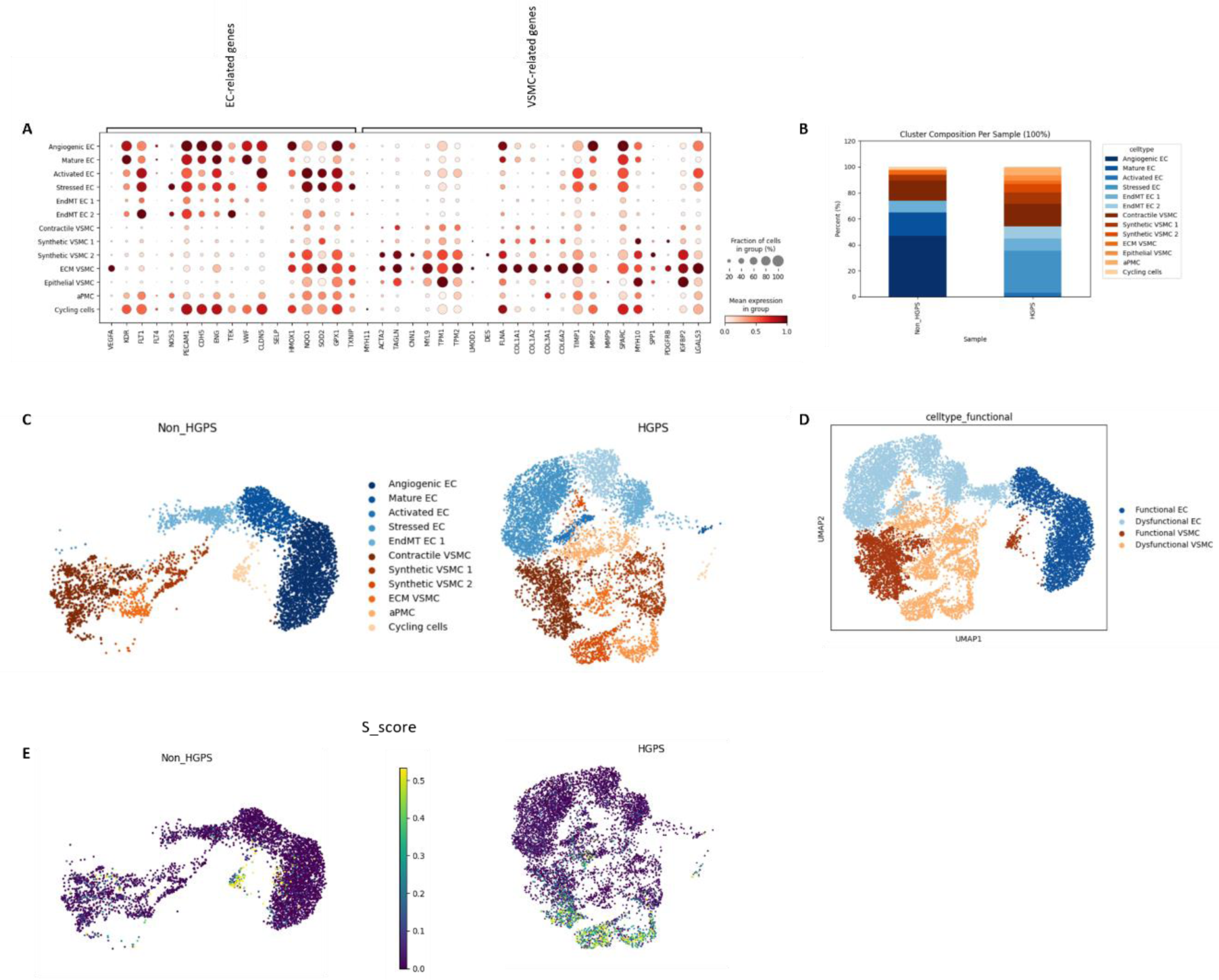
Single cell analysis of accelerated vascular aging in the vascular avatar. **A**, Representative marker genes identified in different ECs and VSMC sub-populations. EndMT EC 1: Endothelial-to-Mesenchymal Transition EC sub-population 1; EndMT EC 2: Endothelial-to-Mesenchymal Transition EC sub-population 2; ECM VSMCs: extracellular matrix VSMCs; aPMC: Activated Mesenchymal (Perivascular Stromal) Cells. **B**, Percentages of different sub-populations of ECs and VSMCs from non-HGPS and HGPS at day 30. **C**, UMAP plots showing ECs and VSMC sub-populations detected in non-HGPS and HGPS vascular avatars at day 30. **D**, UMAP plots showing functional ECs, dysfunctional ECs, functional VSMCs and dysfunctional VSMCs across all cells from non-HGPS and HGPS. **E**, UMAP plots based on cell cycle synthesis scores (S_scores) for non-HGPS and HGPS vascular avatars. A higher S score means that cells have a higher probability of being in S phase.

